# A genomic analysis of parasite-mediated population differentiation in a metapopulation

**DOI:** 10.1101/2022.03.10.483675

**Authors:** Meret Halter, Louis Du Pasquier, Dieter Ebert, Peter D. Fields

## Abstract

Understanding the genetics of host adaptation is a powerful approach to study host – parasite interactions. Hosts are often assumed to have a simple genetic architecture underlying resistance. However, in natural populations the genetics are rarely known and the link between host adaptation and evolutionary models cannot be easily established. To shed light on the genetic basis of host evolution in the presence and absence of parasites we studied a highly dynamic rockpool metapopulation of the planktonic crustacean *Daphnia magna* and its microsporidian parasite *Hamiltosporidium tvaerminnensis*. We examined genome-wide allele frequencies estimated from pooled Illumina sequencing (Pool-seq) of 12 subpopulations of a metapopulation. Subpopulations that had evolved for several years with the parasite were contrasted to uninfected subpopulations, with the aim to find genomic sites of diversifying selection. Consistent with earlier attempts to find resistance genes in this system, we observe many minor-effect outliers, suggesting that the response of the host to this parasite is based on a quantitative-trait architecture. We found 34 outliers across 11 genomic regions that indicate increased differentiation between population groups as well as signs of positive selection. Some of these regions contain immune-related genes, of which four are likely involved in immune downregulation. Our findings show that in the presence of the microsporidium parasite, hosts evolve in a complex polygenic way, driving population differentiation in the metapopulation under study. Such evolutionary differentiation is a powerful mechanism to maintain genetic diversity in spatially structured populations.

## Introduction

Parasites are ubiquitous and affect almost all free-living organisms by limiting growth, altering development, and reducing fecundity and survival of their hosts (Tellier, Brown, Boots, & John). Therefore, parasites and the diseases they cause are major drivers of natural selection (Schmid-Hempel, 2021). Many parasites specialize on one or a few hosts, leading to a tight interaction between the host and its parasite. As a consequence, hosts are selected to reduce the negative fitness effects of the associated disease, for example by evolving resistance. Nonetheless, susceptible hosts can persist in natural populations (Auld et al., 2013; Bento et al., 2020; Henter & Via, 1995). One explanation for the persistence of susceptible hosts in a host – parasite system is that host adaptation comes with a cost, such that resistant individuals are less fit than their susceptible counterparts in the absence of the parasite. Such trade-offs between resistance and other adaptive traits can lead to diversifying selection between populations of the same species that differ in parasite presence (Bartlett et al., 2020; Boëte et al., 2019; Boots & Best, 2018; Leitão et al., 2021). If resistance is costly, one expects that populations without the parasite should lose resistance, while infected populations with maintain or even gain resistance when the parasite is abundant. Consequently, populations with and without the parasite will be under diversifying selection for the genes directly or indirectly associated with the parasite.

Diversifying selection ought to happen even in populations that are geographically close to each other if they are sufficiently isolated and selection is strong enough. A metapopulation system composed of interconnected populations with a co-occurring natural parasite is ideal to study the evolution of the host in the presence and absence of their parasite, because populations, which are otherwise genetically similar, will experience differential selection. In such a case, the populations would show signatures of local adaptation. Therefore, metapopulations may offer new insights into the ongoing process of differential adaption to parasites and to the maintenance of genetic diversity for sites under differential selection (Kawecki & Ebert, 2004).

Allele frequency changes in natural populations depends not only on selection but also on neutral forces like genetic drift and gene flow. While genetic drift stochastically leads to fixation or loss of alleles, gene flow among populations homogenizes local gene pools and reduces differentiation (Holsinger & Weir, 2009). This stochasticity is pronounced in a metapopulation, because the subpopulations are often small and highly dynamic, experiencing local extinction followed by re-colonization of vacant sites (Hanski & Gaggiotti, 2004; Levins, 1970). Thus, metapopulation dynamics differ from dynamics of a panmictic population of the same size: in metapopulations, demographic and stochastic processes, such as genetic drift and founder effects, are important forces shaping the underlying genetic structure (Baguette, Michniewicz, & Stevens, 2017; Feder, Pennings, Hermisson, & Petrov, 2019; Hanski et al., 2017). Directional selection may therefore be less effective within subpopulations (Roulin et al., 2015; Sultan & Spencer, 2002). Nonetheless, in many systems it has been shown that selection leading to local adaptation within metapopulations does happen (Laine, 2008; Peterson, Hilborn, & Hauser, 2014; Thrall et al, 2002), although it is not always found (Roulin et al., 2015).

Metapopulations of the planktonic crustacean *Daphnia magna*, with one of its natural parasites, the microsporidian *Hamiltosporidium tvaerminnensis* (Haag et al., 2011), provide a useful systems to study diversifying adaptation of host resistance (Cabalzar et al., 2019). In the focal metapopulation in South-Western Finland, about 50 % of host subpopulations are infected (Ebert et al., 2001). All genotypes are susceptible, but the hosts show quantitative variation in resistance (Cabalzar et al., 2019; Ebert, 2008). In contrast, in central Europe, complete resistance of *D. magna* to *H. tvaerminnensis* is common (Lange et al., 2015). *D. magna* individuals from the focal metapopulation were shown to adapt rapidly to cope with the parasite without reducing parasite burden (Zbinden et al., 2008), suggesting that resistance to *H. tvaerminnensis* is gained by increasing tolerance towards the parasite. Uninfected populations were shown to evolve tolerance after introduction of the parasite (Zbinden et al., 2008; Bieger & Ebert, 2009; Ebert et al., 2001; Lass & Ebert, 2006). These studies suggest that there is not only positive selection in the presence of the parasite, but also selection against resistance/tolerance alleles in the absence of it. This variation further implies that resistance/tolerance is costly. Indeed, Zbinden et al. (2008) suggested a cost paid in form of reduced asexual growth rate of resistant genotypes.

The genetic basis and mechanism of resistance against *H. tvaerminnensis* is at present not known, but earlier studies identified candidate genomic regions. In these studies, a susceptible *D. magna* genotype from the Finnish metapopulation was crossed with a resistant genotype from Southern Germany (F2-QTL panel analysis) (Krebs et al., 2017; Routtu & Ebert, 2015; Routtu et al., 2014). Eight minor Quantitative Trait Loci (QTL) contributing to resistance were identified. It is, however, unclear if these candidate genomic regions for resistance are of similar importance within the metapopulation in Finland, where all genotypes are susceptible. Here we investigate the genetic structure of resistance to *H. tvaerminnensis* in *D. magna* from the Finnish rock pool metapopulation. To achieve this, we set out to answer the following questions: I) Are there molecular signatures of positive selection in *D. magna* associated with parasitism by its common parasite, *H. tvaerminnensis*, in a metapopulation? II) Are the candidate genetic regions overlapping with the previously described sites for resistance QTLs?

To surmount the difficulties in detecting signatures of positive selection in a highly dynamic metapopulation system, we combine several approaches to identify candidate regions. We study a single natural metapopulation from which we select six subpopulations that have evolved with the parasite and six subpopulations that have evolved without the parasite for several years. We first identify putative outlier SNPs using common outlier-tests. To narrow down regions showing signatures of positive selection, we use population genetic summaries to compare the outlier regions to a neutral genomic background, simultaneously accounting for the specific demography shaping the different summaries. Outliers that show significant difference to the background in statistic summaries are then examined for the genes associated with differentiation. Overall, our results indicate that resistance/tolerance against *H. tvaerminnensis* is most likely a quantitative trait, with many sites of small effect being involved.

## Methods

### Model system

The freshwater crustacean *D. magna* Straus 1820 (Crustacea: Cladocera) has a 4olarctic distribution and is common in rock pools along the coast of the Baltic Sea (Pajunen & Pajunen, 2003). In South-Western Finland *D. magna* inhabits freshwater rock pools of various sizes on Skerry islands, forming a metapopulation structure. The filter-feeding zooplankton is cyclically parthenogenetic and produces primarily asexual daughters, but, following environmental induction, females produce sons to initiate sexual reproduction. The sexually produced resting eggs can endure harsh environmental conditions and hatch only when hatching conditions are good (Ebert, 2005). In spring, females hatch from resting eggs and start to reproduce asexually. The asexual generation time is 10-20 days, and in the focal metapopulation, there are approximately 8-10 asexual generations per year (Zbinden et al., 2008).

All populations used in this study come from a metapopulation system near Tvaerminne Zoological Station, Finland (59°50’N, 23°15’E) that has been monitored since 1982. 565 rock-pools on 16 islands have been checked twice a year for presence of *Daphnia*. From 1998, *D. magna* populations were checked in irregular intervals (but yearly since 2010) for the presence of parasites. Between 40 and 120 of these pools have *D. magna* populations with fluctuations from year to year. The rock pools near Tvaerminne freeze in winter and are at risk of drying up during summer. This constantly changing habitat leads to a high chance of extinction for populations (yearly extinction rates of 17 %). Empty pools can then be re-colonized when conditions change again (Pajunen & Pajunen, 2003). Since dispersal is thought to happen mainly through passive movement of resting stages, aided by wind, water and birds, it is likely that many populations are founded by only a single colonizer (1.8 genotypes on average; Haag et al. 2005).

The most abundant parasite in the Finnish metapopulation is the microsporidian, *H. tvaerminnensis*, formerly referred to as *Octosporea bayeri* (Haag et al., 2011); this parasite also occurs throughout much of the species Eurasian range (Angst et al. 2022). *D. magna* is its only host. It infects host fat cells and ovaries, leading to substantial reduction of host fitness by reducing life span, fecundity, and competitive ability (Ebert et al., 2007; Lass & Ebert, 2006; Vizoso & Ebert, 2004). In later stages of the infection, spores are found in the entire body cavity of the host and are released from decaying cadavers to infect new hosts (Bieger & Ebert, 2009). Besides horizontal transmission, *H. tvaerminnensis* is also vertically transmitted, with 100 % transmission rate to asexual offspring, but generally less than 100 % to sexual offspring (Ebert et al., 2007). The parasite is therefore unlikely to ever be lost from a population once infected (Lass & Ebert, 2006; Vizoso & Ebert, 2005).

### Choice of infected and uninfected populations

For the current study, we chose *D. magna* subpopulations that fulfill three criteria. (i) Infection of *H. tvaerminnensis*: Half of the subpopulations must have been infected for 10 or more years, so that selection by the parasite would have had time to act (approximately 80 to 100 asexual generations, and 10 to 20 sexual generations). The other half of the selected subpopulations must have no present infection since they originated, i.e. new colonizations of empty habitats. This allows us to compare genetic differences between these two natural groups of subpopulations. Age of infection was assessed from the longitudinal parasite checks described above. (ii) Population age: Age is important because over time, immigration reduces the consequence of the strong founder effect during colonization, making older populations on average more genetically diverse and less differentiated from other populations (Walser & Haag, 2012). We therefore included only the oldest populations (youngest infected pop. = 16 yrs; youngest non-infected pop. = 5 yrs). Population age was assessed from the longitudinal time series data (Ebert, et al 2001; Dubart et al., 2020; Pajunen & Pajunen, 2003). A population which was not present for two survey periods (and one winter) in a row is considered extinct. This criterium is supported with genetic data from newly found populations (Haag et al. 2006). (iii) Spatial arrangement of populations: Infected populations included in our study are geographically spread out over several islands, each with at least one uninfected counterpart on the same island. This set up will help to isolate the parasite’s effect on host allele frequencies from other demographic processes, like possible island effects and relatedness.

### Sample collection and pool (re)-sequencing

For each of the focal populations, we collected between a minimum of nine (every adult individual in the population) and up to 59 individual *D. magna* adults in May 2014 after they had freshly hatched from sexually produced resting eggs, thereby ensuring that every female represented a unique genotype. To avoid population extinctions in case we sampled all females, we allowed the females to reproduce in the laboratory and returned the newborns back to their population of origin. To reduce non-focal DNA in our sequencing reactions (from microbiota and food items), individuals were treated for 72 h with a cocktail of three antibiotics (streptomycin, tetracycline, ampicillin) at a concentration of 50 mg/L each, and this treatment was refreshed every 24-hours. Animals were fed with dextran beads (Sephadex ‘Small’ by Sigma Aldrich: 50 μm diameter) at a concentration of 0.5 g/100 mL to aid gut evacuation (Dukić et al. 2016). Animals from each population were moved out of antibiotics and into 2.0-mL Eppendorf microcentrifuge tubes and excess fluids removed with a sterile pipette. RNAlater (ThermoFisher Scientific, catalog # AM7022) was added to each tube in order to fully submerge the animals. This initial aliquot of RNAlater was removed from the tube and discarded. A second aliquot of RNAlater was added to submerge the animals. Each tube was incubated overnight at 4 °C and then subsequently stored at −20 °C until extraction.

In preparation for DNA extraction, RNAlater was removed from each tube with a sterile pipette. The animal tissue was subsequently submerged briefly in purified water in order to dilute the residual RNAlater solution which might inhibit DNA extraction reagents. This mixture was removed and discarded. DNA Extraction buffer (Qiagen GenePure DNA Isolation Kit) was subsequently added to the tubes, and tissue was disrupted using sterile and DNA-free plastic pestle. The resultant solution was incubated overnight with Proteinase K at 55 °C. RNA was degraded using RNAse treatment for one hour at 37 °C. Protein removal and DNA precipitation, including the addition of glycogen (Qiagen) to aid DNA precipitation, were done using the Qiagen GenePure DNA Isolation Kit instructions. Resultant purified DNA was suspended in 40 μL of Qiagen DNA hydration solution and subsequently tested for purity and concentration using a Nanodrop and Qubit 2.0, respectively. Libraries were prepared using Kapa PCR-free kits and sequenced by the Quantitative Genomics Facility service platform at the Department of Biosystem Science and Engineering (D-BSSE, ETH), in Basel, Switzerland, on an Illumina HiSeq 2000 to generate 150 bp paired end (PE) data.

### Variant discovery

Read quality was assessed using FastQC v.0.10.1 (http://www.bioinformatics.babraham.ac.uk/projects/fastqc). Illumina sequencing of the pools of genotypes from the individual populations were adapter and quality trimmed using Trimmomatic v.0.36 (Bolger et al., 2014). After trimming of adapter sequences, terminal bases with a quality score below three were removed from both ends of each read. Then, using the sliding window function and again moving in from both sides, further 4 bp-fragments were removed if their average quality score was below 15. Read quality was re-checked with FastQC to confirm quality and adapter trimming succeeded. Retained reads were mapped to the *D. magna* 3.0 reference genome (Ho et al., 2020; P. Fields et al. In Prep.; BioProject ID PRJNA624896) using BWA MEM (v. 0.7.13) (Li, 2013; Li & Durbin, 2009). Importantly, this reference genome derives from a genotype collected from a rock pool near the Tvaerminne field station and is therefore closely related to the samples used in this study. In addition to default parameters, we added the Picard comparability (-M) option (Li & Durbin, 2009). The resulting SAM files were subsequently converted to BAM files, coordinate sorted, and filtered for mapping quality ≥ 20 using SAMtools version 1.3 (Li et al., 2009). Individual bam files were given a read-group name with Picard Tool’s AddOrReplaceReadGroups (Picard Toolkit, 2019) and reads were realigned around indels with IndelRealigner from the Genome Analysis Toolkit (GATK) version 3.7 (McKenna et al., 2010).

Variant calling for the individual population samples and the pooled infected and uninfected population samples was performed with the UnifiedGenotyper from GATK (DePristo et al., 2011), excluding reads with a mapping quality less than 30 (see Suppl. Figure 1 for a summary of our pool-sequencing approach for variant discovery). Further filtering criteria included only biallelic SNPs with a quality-score > 30, a mapping quality > 40, quality by depth > 2 and Fischer strand < 60, following the GATK recommendations (Van der Auwera et al., 2013). A final filtering step was added for the individual population samples: (i) including only sites for which all populations had more than 15 reads and (ii) the minor allele was represented by at least two reads in at least two different populations. Prior to SNP calling for the pooled samples, we first made sure that all populations were represented with equal coverage in the pooled samples by down sampling individual population bam files with high coverage to the coverage of the population with lowest mean coverage of each pool. Coverage analysis and down sampling were done with SAMtools coverage, v. 1.3 (Li et al., 2009) and Picard Tool’s Downsample (Picard Toolkit, 2019) with a random down-sampling algorithm. The down-sampled bam files were then merged with SAMtools merge, and SNP calling was performed in the same manner for individual population samples. The final filtering steps for the pooled samples consisted of (i) filtering for read counts greater than 60 (6 populations x 15 reads) and (ii) considering only sites with a minor allele count larger than 4 (2 populations x 2 minor alleles).

### Genome scan for adaptive differentiation

To detect genomic signatures of divergent selection we relied on two approaches. The R package PCAdapt (v. 4.0.3; Luu et al., 2017) identifies loci as outliers when they are excessively related with population structure, as assessed by a Principal Component Analysis (PCA). When using the individual populations as separate inputs for the PCAdapt package, we observed an inflation of very low p-values. PCAdapt was run with the default settings for Pool-seq data. Multiple testing p-value inflation was controlled for using the R package qvalues, v. 3.7 (Storey et al., 2018) with ***α*** set to 0.1.

The second software package used to detect genomic signatures of selection was BayPass v. 2.1 (Gautier, 2015). BayPass first identifies overly differentiated SNPs based on the *XtX* statistic and in a second step identifies SNPs associated with a population-specific covariable. The implemented framework of BayPass explicitly accounts for population structure originating from shared history of the focal populations with a covariance matrix. Therefore, BayPass is less sensitive to the confounding effects of demography and selection. The core model was run both with the pooled allele count data and the individual population allele count data. All parameters were set to default, except for the number of post burn-in and thinning samples set to 10,000 for the Markov Chain Monte Carlo (MCMC) options. Under the core model the *XtX* was estimated for each SNP as well as the covariable matrix Ω for the examined populations. A pseudo-observed data set (POD) simulated with the same parameter values as the real data were used to define a threshold for identifying overly differentiated SNPs.

In a second step, the association of allele frequencies to infection status was tested with the auxiliary variable model (AUX) for each individual population and with the less complex standard covariate model (STD) for the pooled allele count data. In the AUX model, a Bayesian binary auxiliary variable is introduced to indicate whether a specific SNP can be regarded as associated with the environmental covariable (infected/non-infected) or not. The Bayes Factor (BF), expressed in deciban units (dB) is then derived from the posterior probability of association for each SNP, while accounting for multiple testing issues. We followed the decision criterion of Jeffreys (Jeffreys 1961) to quantify the strength (in dB) of evidence of the association of a given SNP to the covariable. Jeffrey’s rule states “very strong evidence” when 15 dB < BF < 20 dB and “decisive evidence” when BF > 20 dB. The STD model that we used for the pooled allele count data evaluates a linear association to the trait of interest with the approximate Bayesian p-value in the log10 scale (eBPmc), and therefore the decision criterion for associated SNPs is less straightforward.

Each model was run three times with a different initial seed of the (pseudo-)random number generator. The posterior means of all three iterations were found to be very similar with the Ω matrix having a FMD distance < 0.01 between each pair (Förstner & Moonen, 2003). We only considered a SNP for further analysis when it exhibited an *XtX* value > 1 % POD and BFmc > 20 dB or an eBFmc < than the 0.001 % quintile in all three independent MCMC runs.

### Population genetic summary of outlier regions

To further our understanding of how selection related to parasite presence may have shaped genetic diversity and the site frequency distribution (SFS) of the outlier regions, we examined a 12,000 bp (12 kbp; ± 6 kb) flanking region centered on each of the outliers associated with infection (different window sizes and a bigger flanking region lead to essentially the same results, see Suppl. Table 1). For each 12 kbp region, we calculated the number of segregating sites per site, **θ_w_** (Watterson, 1975) and the average pairwise difference per site, **θ_π_** (Nei & Li, 1979). Additionally, we calculated the difference between the two diversity estimates, or Tajima’s *D* (Tajima, 1989). Finally, Fay and Wu’s *H* (Fay & Wu, 2000) was calculated, a statistical test that incorporates data from an outgroup species in order to distinguish between the ancestral and derived state of the polymorphism. We used short-read data derived from *D. similis* (NCBI BioProject PRJNA744861; Fields et al. 2022) aligned to the D. magna 3.0 reference genome to polarize variants as part of Fay and Wu’s *H*. Both statistics — Tajima’s *D* and Fay & Wu’s *H*—test the hypothesis that the region experienced recent positive selection, since low and high frequency alleles increase under such a scenario, compared to neutrality (Pavlidis & Alachiotis, 2017). All summaries were calculated with NPStat (v. 3) (Ferretti et al., 2013), a program explicitly designed for pool-seq data (Ferretti et al., 2013). Additionally, we calculated an *F*_ST_ related value specifying the difference between groups (the pooled infected and the pooled non-infected populations) in allele frequencies to the within group. To avoid confusion, we will refer to our *F*_ST_ related value as *F*_GT_ (G for group). Calculation was done with popoolation2 (Kofler et al., 2011) with the default settings for pool-seq data.

We compared all summaries (***θ_w_***, ***θ_π_***, Tajima’s *D*, Fay & Wu’s *H* and *F*_GT_) of the flanking regions of each outlier SNP to a genome-wide sample of flanking regions of 10,000 randomly drawn reference SNPs using a Wilcoxon signed-rank test for differences in the two samples. To account for the differences in the infected and non-infected populations, we obtained the mean of all infected and respectively all non-infected individual population samples and tested them against their neutral background. The basis of this comparison means each outlier-flanking region could either be significantly different from the genetic background in the non-infected populations, the infected populations, or all populations. Only outliers that exhibited a significant decrease in their flanking region of Fay & Wu’s *H*, genetic diversity ***θ_w_*** and ***θ_π_***, and Tajima’s *D* as well as a significantly higher *F*_GT_ than the randomly drawn SNPs were kept for further analysis of their environment.

In addition to the summary statistics of the individual populations, we calculated all summaries for the infected and the non-infected pooled samples as well as a pooled samples containing all populations, which we called the pooled meta sample (aligning and down sampling was done the same way as we did for the other two pooled samples. For the pooled samples, we wanted to test whether we could find flanking regions of outliers that show reduced diversity (i.e., ***θ_w_***) in the infected or in the non-infected pooled samples but showed higher diversity in the pooled meta sample at the same time. The pooled meta sample is expected to have a higher diversity if the infected and non-infected samples are differentiated at a given locus. Since the pooled samples contain less diversity, because of the necessity of down sampling, and with the *F*_GT_ value we include a similar but more accepted way of measuring within-group variability against between-group variability, we decided to not include this summary as a filtering criterion for candidate sites.

### Functional annotation of outlier and population genetic summaries

For coding regions intersecting the 12 kbp flanking regions around each outlier SNP with confirmed signs of positive selection (candidate SNPs) on either the infected or the non-infected populations, we calculated nucleotide diversity (***π***), and its nonsynonymous and synonymous partitions (***π_N_*** and ***π_S_***, respectively), as well as mean nonsynonymous and synonymous divergence (***d_N_*** and ***d_S_***) from the reference sequence. This basic comparison allows us to, theoretically, distinguish positive selection from purifying selection or random genetic drift: ***π_N_*** = ***π_S_*** indicates neutrality, ***π_N_*** < ***π_S_*** indicates purifying selection, and ***π_N_*** > ***π_S_*** may indicate positive selection at protein coding sites (Hughes, 1999). To account for demographic effects, we compared the mean ***π_N_/π_S_*** ratio of the genome-wide sample of genes in the flanking regions of 10,000 randomly drawn reference SNPs to the genes in the flanking regions of each outlier using a Wilcoxon signed-rank test. We identified coding regions intersecting the 12 kbp flanking regions with BEDtools (v. 2.25.0 Quinlan & Hall, 2010). We used SNPGenie (v.3; Nelson et al., 2015) to generate estimates of nucleotide diversity and genetic distances. Finally, for individual outlier regions we predicted amino acid sequences (Burge & Karlin, 1997) and used BLAST to compare these putative protein sequences against the NCBI protein database (States & Gish, 1994). For genes with predicted functions and gene families in regions around candidate SNPs, we attempted to identify and functionally describe known immunological roles of these gene families (see Suppl. Figure 2 for a summary of our outlier and population genetic summary statistic approach).

As a final step in estimating the strength of directional selection on the outlier SNP loci vs. the genomic background, we used SelEstim v.1.1.7 (Vitalis et al., 2014). We focused primarily on the pooled allele counts of infected vs. non-infected populations, because the results of individual populations and pooled allele counts were very similar. We relied on primarily default parametrizations of SelEstim, and five pilot runs of 1,000 iterations each, followed by 100,000 iterations of burn-in, and then 50,000 iterations of sampling every 20 steps. SelEstim estimates a locus-specific selection coefficient ***δ_j_***. It was used as a measure of the strength of selection, as well as generating a Kullback-Leibler divergence (KLD) to measure the distance of the posterior distribution of ***δ_j_*** from the centering distribution (Vitalis et al., 2014). We generated a simulated POD with the same parameter values, as was done with BayPass above, to define a threshold for identifying SNPs experiencing directional selection based on the KLD. In addition to the basic functionality of SelEstim for determining if particular loci show evidence of directional selection, we also directly compared the distribution of KLD values of previously identified outlier loci and the genomic background using a Wilcoxon rank test.

## Results

### Choice of infected and uninfected populations

Ours election criteria resulted in a total of 12 subpopulations—six infected and six uninfected—spread over five islands (Table 1; Figure 1). Mean population age was 22.5 years and mean genetic diversity of all populations was ***θ_w_*** = 0.0048. Both in age and genetic diversity, the six infected populations exhibit higher (but not significant) means, with a mean age of 26.5 years and mean ***θ_w_*** = 0.0051 as compared to 18.3 years and *θ_w_* 0.0045 of non-infected populations (t-test; p-value for age = 0.09, p-value for ***θ_w_*** = 0.28). This is consistent with the earlier observation that longer lasting populations have a higher probability of acquiring the parasite (Ebert et al., 2001) and that infected populations tend to have higher genetic diversity (Cabalzar et al., 2019).

**Figure 1.**
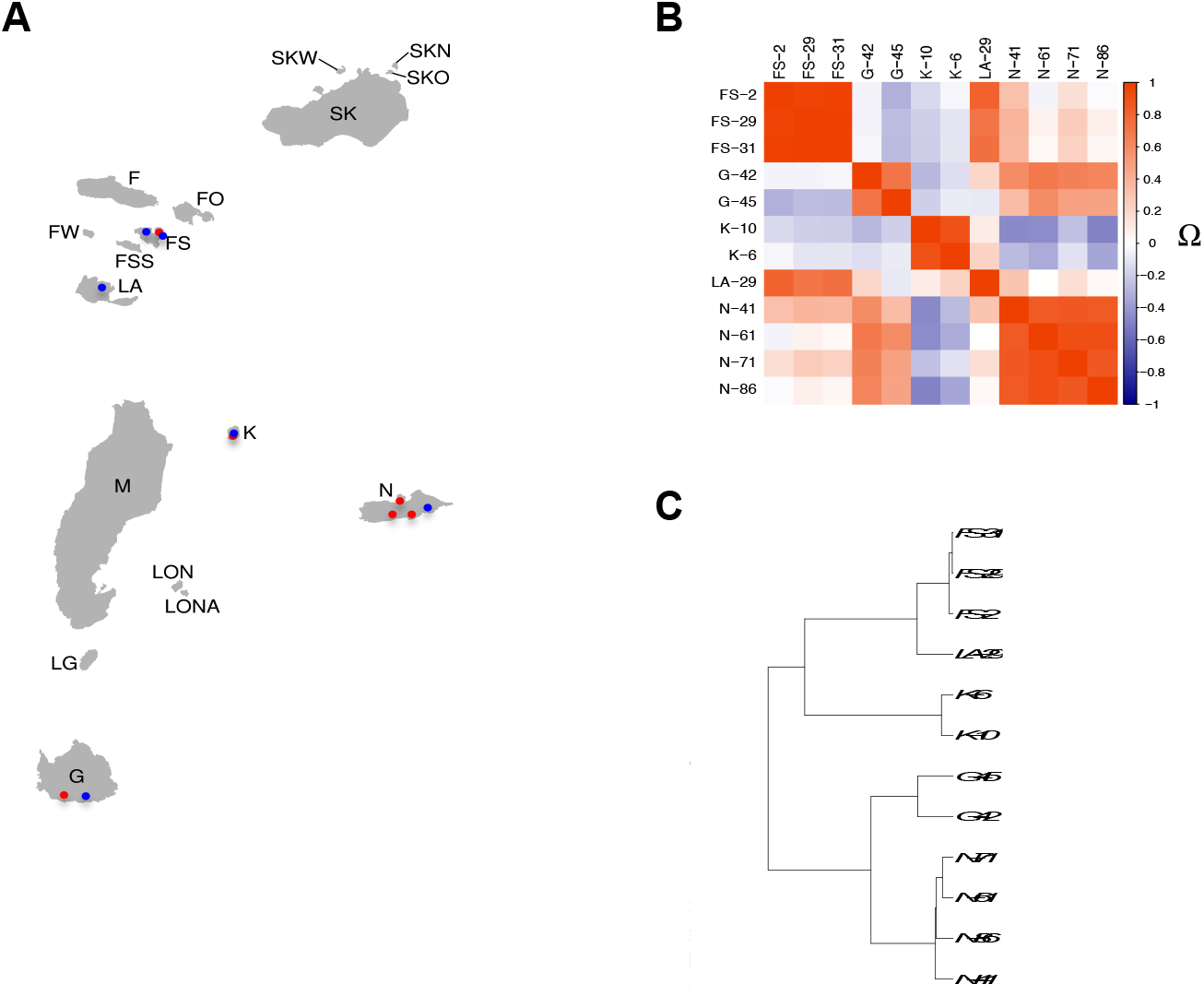
Samples used to identify signatures of divergent selection arising from host-parasite coevolution. A) Sampling area of the long term metapopulation project. locations in the Finnish metapopulation of *D. magna*, where red dots indicate samples of populations infected by *H. tvaerminnensis* and blue does indicate samples of populations not infected; B) Correlation plot and C) hierarchical clustering dendrogram derived from the Ω – matrix resulting from the core model of BayPass.

**Table 1.**
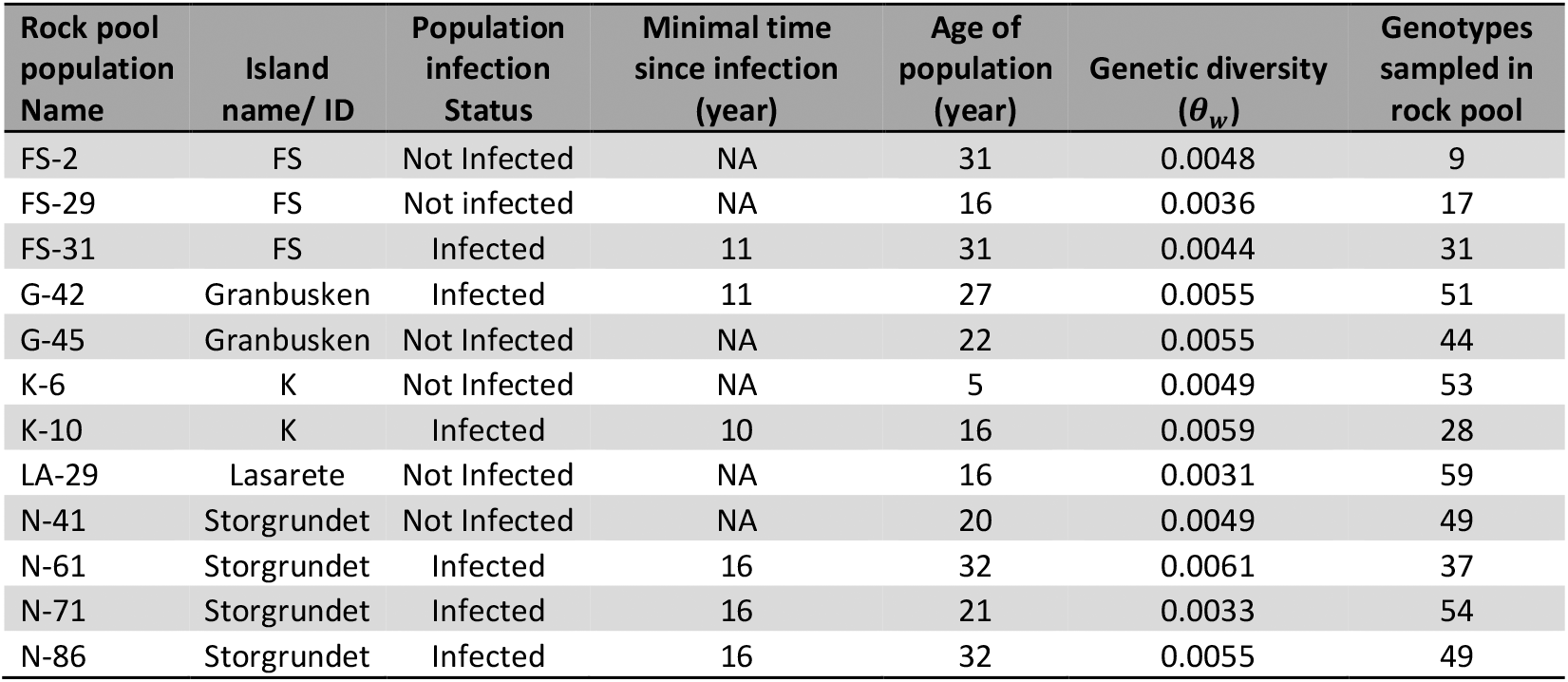
Summary of the 12 samples from the focal metapopulation in the Tvärminne archipelago in Southwestern Finland. Nameless islands are called by a letter ID.

| Rock pool<br>population<br>Name | Island<br>name/ ID | Population<br>infection<br>Status | Minimal time<br>since infection<br>(year) | Age of<br>population<br>(year) | Genetic diversity<br>($\theta_w$) | Genotypes<br>sampled in<br>rock pool |
| --- | --- | --- | --- | --- | --- | --- |
| FS-2 | FS | Not Infected | NA | 31 | 0.0048 | 9 |
| FS-29 | FS | Not infected | NA | 16 | 0.0036 | 17 |
| FS-31 | FS | Infected | 11 | 31 | 0.0044 | 31 |
| G-42 | Granbusken | Infected | 11 | 27 | 0.0055 | 51 |
| G-45 | Granbusken | Not Infected | NA | 22 | 0.0055 | 44 |
| K-6 | K | Not Infected | NA | 5 | 0.0049 | 53 |
| K-10 | K | Infected | 10 | 16 | 0.0059 | 28 |
| LA-29 | Lasarete | Not Infected | NA | 16 | 0.0031 | 59 |
| N-41 | Storgrundet | Not Infected | NA | 20 | 0.0049 | 49 |
| N-61 | Storgrundet | Infected | 16 | 32 | 0.0061 | 37 |
| N-71 | Storgrundet | Infected | 16 | 21 | 0.0033 | 54 |
| N-86 | Storgrundet | Infected | 16 | 32 | 0.0055 | 49 |

### Variant discovery

Sequencing of all 12 populations produced 884 million reads with an average of 74 million reads per sample. Of these reads, 94.95 % mapped to the *D. magna* reference and 90.28% were properly paired. Duplicate filtering removed 4 % of all mapped reads. After indel realignment and duplicate removal, remaining reads covered 99.97 % of the reference with an average depth of 58× per sample. When we used individual populations as the input for variant calling, we obtained in total 1,894,780 SNP variants. Quality filtering reduced SNP number to 1,165,194 variants, and insufficient coverage among populations reduced the number of retained variants to 815,338. These SNPs were used in downstream analysis, giving us ^~^ 5.8 SNPs per kbp. For the pooled population groups (infected and non-infected) we obtained a total of 1,249,996 SNP variants. Quality filtering reduced the SNP number to 711,408, and removal of SNPs with insufficient coverage for one of the population groups lead to the final number of 673,876 SNP variants used for further analysis (^~^ 4.8 SNPs per kbp). The difference in number of SNP variants between the two pooling strategies resulted primarily from the down sampling approach, which evens out the differences in coverage among populations.

### Genome scans for adaptive differentiation

Under the core model of BayPass, we estimated the scaled covariance matrix **Ω** for the 12 individual population allele frequencies, quantifying the genetic relationship among all pairs of populations. The estimates of **Ω** seem to generally reflect a signal of isolation-by-distance (IBD), where populations sampled from the same island generally showing a larger co-ancestry coefficient (Figure 1). A POD sample containing 800,000 SNPs was used to calibrate the *XtX* for each SNP. 5022 SNPs were identified as overly differentiated at the significance threshold of 1% POD. Under the auxiliary covariate model, 7,384 SNPs showed a BF > 20. However, only 484 SNPs of the *XtX* outliers also showed a BF > 20.

The BayPass core model was also used to analyze the pooled infected and non-infected samples. The auxiliary covariate model failed to reach satisfactory convergence, possibly due to specific expectations of the BayPass approach, including the inclusion of more than two populations. The results of the covariate model for association of the infection status were identical to the *XtX*, which is also expected since the infection status splits these two populations, i.e., the difference of infection status is identical to the difference between the two populations. We therefore used the 0.001 quantile as a stringent significance threshold (0.1%), resulting in 244 of the most differentiated SNPs between the two samples.

With the pooled infected and non-infected samples, PCAdapt found 31 outliers among the ^~^ 670,000 SNPs. These 31 loci exhibited the most significant population structure when adjusting for multiple testing with q-values (alpha = 0.1). These three different outlier-scans resulted in 705 unique outlier SNPs (51 were found both in the individual BayPass analysis as well as the pooled BayPass analysis). An overlap between outliers of all three scans could only be seen in two regions of about 100 bp in length (contig 04F from 3191160-3191243 bp; linkage group 2; and on contig 11F 1389176-1389280 bp; linkage group 4; Suppl. Figure 4).

### Population genetic investigation of outliers

When comparing the population genetic summary statistics (***θ_w_***, Tajima’s *D*, Fay and Wu’s *H*, and *F*_GT_) of individual populations, each summary differed significantly between outlier plus flanking (12 kbp) regions and the background genomic regions. Overall, for the 705 outliers, ***θ_w_***, Tajima’s *D* and Fay and Wu’s *H* were decreased compared to background, while *F*_GT_ between the pooled populations was increased (Table 2, all populations). The reduced diversity in outliers is also reflected in the genic polymorphisms summarized in ***π_N_***, ***π_S_***, ***d_N_***, ***d_S_***, and ***π_N_/π_S_***. Both ***π_N_*** and ***π_s_***, as well as ***d_N_*** and ***d_S_***, are reduced at outlier + flanking region SNPs as compared to the genomic background (Table 2). Finally, the observed ***π_N_/π_S_*** of the outliers + flanking region is generally reduced compared to the background (Table 2).

**Table 2.**
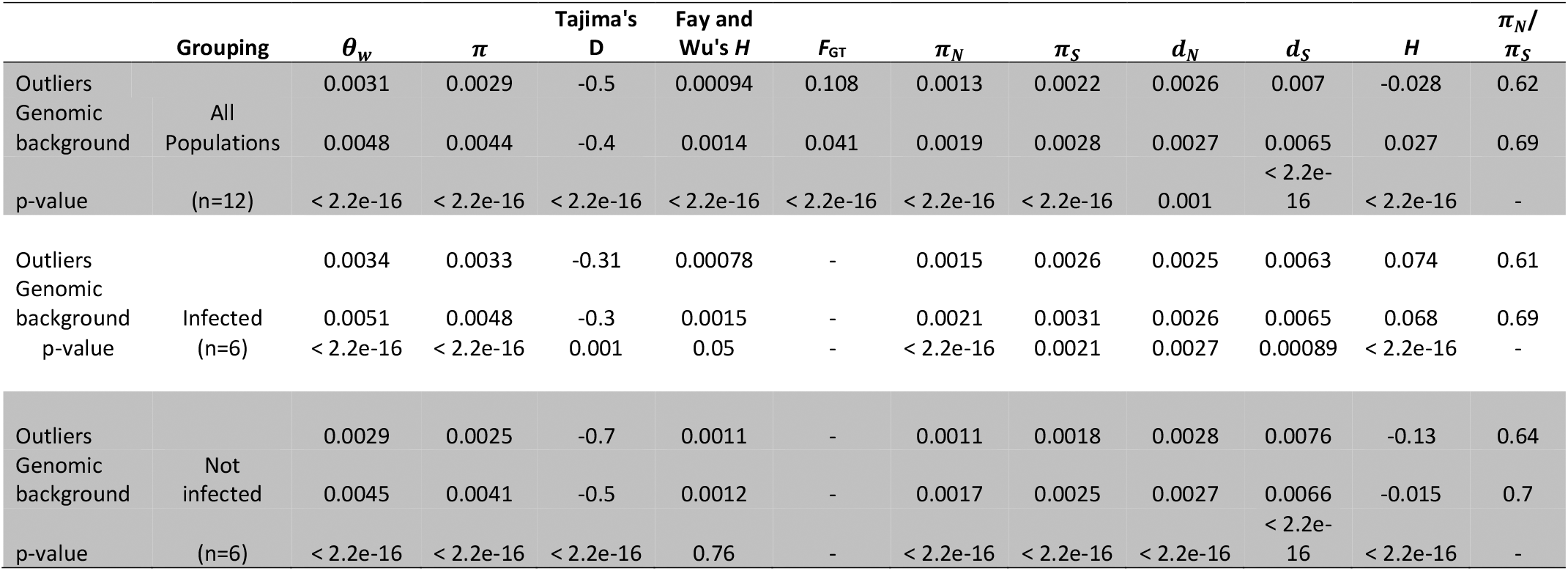
Summary statistics for both genic and non-genic regions at outlier loci + flanking regions and for the genomic background. Summaries of all 705 outlier SNPs + flanking regions and the flanking regions of randomly drawn SNPs. P-values of the Wilcoxon rank test between the two groups of each summary are reported. ***π_N_*** and ***π_S_*** as well as ***d_N_*** and ***d_S_*** are calculated only for polymorphisms inside genes intersecting the flanking regions. *H* is a measure of gene diversity or expected heterozygosity.

When we distinguish between infected populations and non-infected populations, for both outlier and background sites, we observe an increased diversity in infected populations (Table 1 and 2). Infected populations also have more polymorphisms in coding regions than the non-infected populations, both in outliers and background flanking region. The same trends between the outliers and the background can also be seen in the two population groups, e.g., Fay and Wu’s *H* (Table 2). Out of the 705 outliers + flanking regions, only 34 SNPs showed significant deviance from the background sample in all four summary statistics (***θ_w_***, Tajima’s *D*, Fay & Wu’s H and *F*_GT_). These 34 candidates group into 13 regions on nine contigs and six linkage groups.

Mean selection coefficients, where larger values indicate stronger selection, as measured by SelEstim, were 0.35 (std. dev. 0.36) for the whole genome and 0.97 (std. dev. 0.62) for the outlier loci (Figure 2). We found a significant difference (*p* < 2.2e-16) between the distribution of selection coefficients for the genome wide variants and the outlier loci with a Wilcoxon rank test (Figure 2).

**Figure 2.**
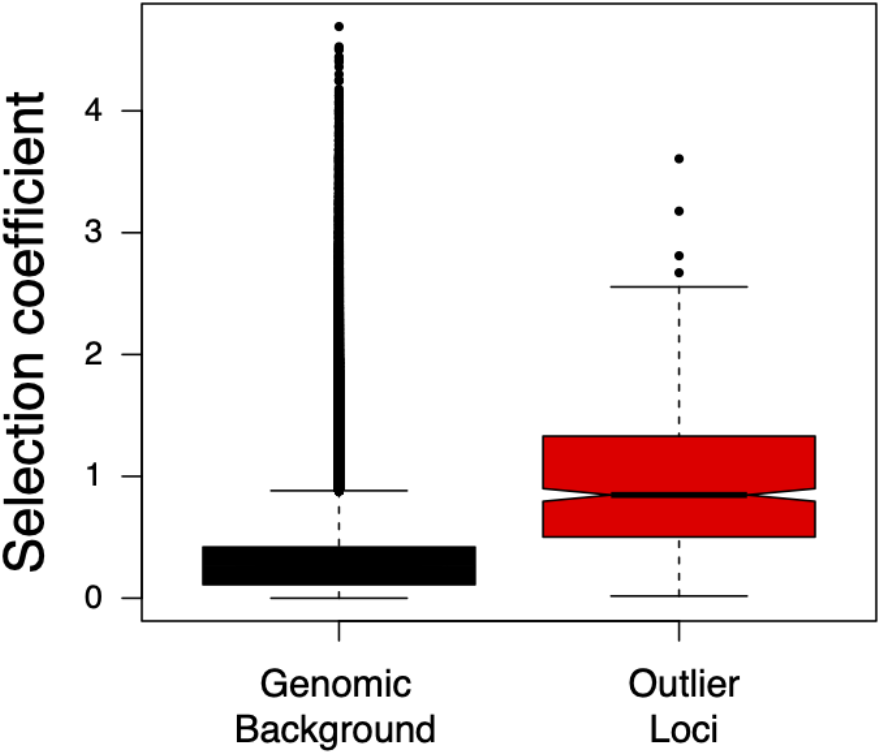
Selection coefficients estimated with SelEstim for all variants and for loci identified with the BayPass approach. Outlier loci exhibit a mean selection coefficient which is significantly greater (almost 3X) than the genomic background (*p* < 2.2e-16).

### Functional annotation of outlier and population genetic summaries

We found 24 genes within the 12-kbp flanking the 34 candidate SNPs, which were outliers across multiple different outlier detection approaches, with about 1.5 genes per 12-kbp (Table 3). In contrast, the gene count of all outlier + flanking regions is 2.9 genes per 12 kbp and 3.37 genes per 12 kbp in the background flanking regions. There were eight candidate SNPs which occurred within a gene (coding sequence or intron), and, out of the 24 genes, five had an elevated ***π_N_/π_s_*** ratio in the infected or the non-infected populations. Five more genes showed only changes at non-synonymous sites. 14 genes showed high similarity to gene families with known function in a non-*Daphnia* model species, of which five gene families have been shown to play a role in immune regulation in other animals and another two genes belong to prominent transporter gene families without being direct immunity-related genes, transporter in immune cells regulate their activity (e.g., control of access to nutrients) and are vital factors in the immune responses. Two genes have uncertain functional predictions of a putative domain or gene family, and nine genes are recognized, but with unknown putative function. These genes are only found in *D. magna* and close relatives.

**Table 3.** Characteristics of the 13 candidate sites. Regions are specified by contig name; for several regions on the same contig subscripts A, B and C specify the order on the contig. ^1^ = Immune gene; ^2^ = transporter gene(without being direct immunity-related genes transporter in immune cells regulate their activity (control of access to nutrients e.g.)and are vital factors in the mmu e responses.;; ^3^ = candidate region overlaps with previous QTL analyses; ^4^ = candidate gene inferred to be part of immune gene cluster.

| Contig | Linkage group | Putative gene annotation |
| --- | --- | --- |
| 08F<sub>A</sub><sup>4</sup> | 1 | All identified genes have no known function |
| 08F<sub>B</sub><sup>4</sup> | 1 | Smyd (=SET and MYND domain-containing protein 4)<sup>1</sup> (down regulation) and MFT (=Major facilitator superfamily transporter)<sup>2</sup> |
| 08F<sub>C</sub><sup>4</sup> | 1 | GT (=Beta-1,4-glucuronyltransferase 1)<sup>1</sup> |
| 09F | 3 | Identified genes with either no know function or non-immune related functions |
| 11F<sup>3,4</sup> | 4 | MORG1 (=Mitogen- activated protein kinase organizer )<sup>1</sup> |
| 23F<sup>3</sup> | 4 | F-box (=F-box/LRR-repeat protein 14)<sup>1</sup> (down regulation) |
| 26F<sub>A</sub><sup>3</sup> | 4 | ABCH (=ABC transporter subfamily H.)<sup>2</sup> |
| 26F<sub>B</sub><sup>3</sup> | 4 | No putative genes identified |
| 01F | 5 | All identified genes have no known function |
| 05F<sup>3</sup> | 8 | Non-immunity related genes |
| 14F<sub>A</sub><sup>4</sup> | 9 | Non-immunity related genes |
| 14F<sub>B</sub> | 9 | Identified genes with either no known function or non-immune related functions |
| 59F | 9 | DGK (=Diacylglycerol-kinase beta)<sup>1</sup> (down regulation) |

The 24 genes showing distinct outlier status across the different applied approaches are unevenly distributed across the 13 candidate regions (Table 3). One region exhibits no genes in the 12-kb kb flanking region. In two regions, only non-immune-related genes are found that exhibit no difference in ***π_N_*** or ***π_s_*** (contig 05F and 14F) as compared to the genome background. In four regions, the genes exhibiting an elevated ***π_N_/π_s_*** rate are uncharacterized or of uncertain functional annotation. In two regions, there are two immune-related genes which appear to be experiencing purifying selection (elevated ***π_s_*** versus ***π_N_***). And finally, four regions have immune-related genes or transporter genes with an elevated ***π_N_/π_s_*** ratio.

## Discussion

In subdivided populations, spatial heterogeneity can maintain genetic variation at loci under differential selection. In the metapopulation studied here, about half of the populations harbor a virulent microsporidian parasite, presumably driving hosts to evolve resistance and tolerance. On the other hand, resistance/tolerance has been shown to be costly in this system (Zbinden et al., 2008), presumably driving parasite-free host populations to lose their adaptation to the parasite. This scenario offers us the opportunity to search for genomic regions with differential selection associated with parasitism. To find sites of differential selection in the metapopulation due to *H. tvaerminnensis*, we compared allele frequencies amongst six infected populations and six non-infected populations. A combination of genome-wide approaches revealed 705 SNPs putatively associated with parasite presence. We then tested the outlier flanking regions for genetic signatures of positive selection using common population genetic summaries, leaving us with 34 candidate SNPs, which are located in 13 well-defined regions on 9 linkage groups. Of these 13 regions, 5 coincide in location with previously described QTLs (Krebs et al., 2017; Routtu & Ebert, 2015; Routtu et al., 2014). We found that, out of the 24 genes within the flanking region of the candidate SNPs, five genes are known for their role in immune regulation in other systems, of which four genes potentially down-regulate the host’s immune function. Finally, we show that the strength of selection taking place on these outlier regions is nearly three times as strong as compared to the genomic background.

Taken together, our study revealed several regions where genetic variation for host-parasite interactions can be maintained by differential selection. Furthermore, our results are consistent with previous findings about a polygenic relationship of *D. magna* and *H. tvaerminnensis*, with 13 putative regions under positive selection in parasitized populations in the metapopulation playing a role in host adaptation. The same variants are presumably selected against in the parasite-free populations. The overlap between some of the previously found QTLs for host resistance and the candidate SNPs of the present study strengthens the importance of these regions for host defense to *H. tvaerminnensis* (Suppl. Table 2). With the finding of five genes involved in immune regulation at candidate regions, we have found promising candidates underlying diversifying selection in the metapopulation. An obvious conclusion is that the trait underlying interactions between *D. magna* and *H. tvaerminnensis* is polygenic in nature.

### Outlier approach and signs for positive selection

Since our natural populations come from a highly dynamic metapopulation, we decided to use two outlier analysis tools and two pooling approaches for our analysis. This way we screened the genome for enhanced population structure and enhanced differentiation between groups with diminishing differences among islands. We were also able to test for an association with the infection status in individual populations. Outlier tests, however, are prone to high false positive rates (François et al. 2016), which is why we used population genetic summary statistics (Garud et al., 2015; Yoder et al., 2014) and *F*_GT_ (Holsinger & Weir, 2009) to compare outliers to the neutral genetic background of the metapopulation. Hence, we were able to verify whether outlier genetic signatures are conforming to our expectation of differential selection acting in the metapopulation.

The summaries of our outlier SNPs can be best explained under a scenario of positive selection. Outlier flanking regions have low genetic diversity (***θ_w_***) and an excess in both low and high allele frequencies (Tajima’s *D* and Fay & Wu’s *H*) when compared to the neutral genetic background. Low genetic diversity and excess in low and high allele frequencies are suggestive of positive selection acting on the outlier SNPs (Braverman et al., 1995; Fay & Wu, 2000; Maynard Smith & Haigh, 2007). This is consistent with the idea that adaptation to *H. tvaerminnensis* might happen by hosts evolving tolerance, which is thought to be subject to positive selection. We also find elevated *F*_GT_ between population groups compared to the neutral genetic background (Table 2), which is consistent with differential selection in the different population groups. Additionally, when we compare the selection coefficient as estimated with SelEstim, we find the outlier loci appear to be experiencing a selective strength nearly 3× higher than the genomic background (Figure 2), and some of the highest selection coefficients in the entire genome (Suppl. Figure 3). Interestingly, while selection coefficients were distinct between outlier and non-outlier loci, the absolute magnitude of the selection coefficients were quite low when compared to other studies which have applied this same method (e.g. see Walden et al., 2020). Several previous studies have suggested that the efficacy of selection may be reduced in the focal metapopulation (Fields et al., 2018; Roulin et al., 2015).

Genetic variation for resistance and tolerance has been reported in this system (Ebert, 2008; Zbinden et al., 2008) as well as variation in parasite adaptation (Altermatt & Ebert, 2008). Variation in host defense is expected when the host s degree of resistance is determined by host cell immune response, as has been discussed for *D. magna* resistance to *H. tvaerminnensis* (Routtu & Ebert, 2015). There seems to exist no barrier defense against microsporidians (Wittner & Weiss, 1999). It is therefore not entirely unexpected that, here and in earlier studies on this system, a complicated, quantitative trait architecture underlying host resistance, with several minor loci contributing to the phenotype, was observed (Krebs et al., 2017; Routtu & Ebert, 2015). Our current study is consistent with this. We find 13 regions under positive selection distributed across the whole genome. We agree with the earlier suggestion that host resistance is a quantitative trait with several regions in the genome contributing to it.

### Overlap with QTLs and immune gene clusters

So far nine QTLs for resistance of *D. magna* to *H. tvaerminnensis* have been described (Krebs et al., 2017; Routtu & Ebert, 2015). With this analysis we found five regions in proximity of four of these QTLs (Table 3 & see supplementary Table 1). It is not surprising that we did not find an overlap with all QTLs, since the earlier identified QTLs were identified from a F2 panel composed of a resistant *D. magna* clone from Southern Germany and a susceptible/tolerant Finnish *D. magna* clone (Routtu & Ebert, 2015). It is therefore likely that the other five QTLs that were identified in the QTL analysis, highlight important regions for resistance rather than tolerance.

A genome-wide search for genes, implicated in some form of immune-related function in other animal genomes resulted in the description of several well-defined genomic clusters in the *D. magna* genome (Du Pasquier et al. In Prep.). These gene clusters (in the following immune clusters) contain homologs of genes involved in the immune systems of invertebrates and vertebrates (such as complement components, lectins, MyD88, Beta 1,3 GR, Tachylectin, Hematopoetic transcription factors) that segregate together. Five of our nine candidate regions fall into such immune clusters (Table 3). This highlights the contribution of several sites and genes for the defense of *D. magna* to *H. tvaerminnensis*. Furthermore, it suggests that genes involved in immune function against *H. tvaerminnensis* may also play a role against other parasites of *D. magna*. Finally, the same genes may also play a role fighting against infectious diseases in other organisms.

Out of the 13 candidate regions that are suggestive of positive selection, six regions harbor either a gene with a known role in immune regulation (five genes) or a transporter gene (two genes) (Table 3 and 4). In four candidate regions, those genes have an elevated ***π_N_/π_s_*** ratio, suggestive of positive selection.

**Table 4.** Coding sites, name and function of all candidate proteins. All proteins and their putative function of the 13 candidate regions. Match completeness describes whether the whole or only a partial match of the annotated protein to the NCBI RefSeq database was found; candidate SNP suggests whether the focal SNP was found inside the coding sequence or not; elevated ***π_N_*** vs. ***π_S_*** indicates if a particular variant region showed an elevated ***π_N_/π_S_*** ratio.

| Contig | Start Pos. | End Pos. | Protein name | Match Completeness | Putative Function | Candidate SNP | Elevated $\pi_N$ vs. $\pi_S$ |
| --- | --- | --- | --- | --- | --- | --- | --- |
| 01F | 1127098 | 1140978 | Uncharacterized protein | whole | Unknown | yes | yes |
| 05F | 2722133 | 2722658 | Protein Z-dependent protease inhibitor | partial | Inhibits activity of the coagulation protease factor Xa in the presence of PROZ, calcium and phospholipids | no | no |
| 05F | 2721163 | 27373700 | Phorbol ester/diacylglycerol-binding protein unc-13 | whole | May form part of a signal transduction pathway, transducing the signal from diacylglycerol to effector functions. Involved in many possible processes | yes | no |
| 08F | 535067 | 538371 | pre-mRNA-processing factor 39 | partial | Involved in pre-mRNA splicing | yes | no |
| 08F | 544162 | 555534 | Uncharacterized protein | whole | Unknown | no | no |
| 08F | 779226 | 779557 | SET and MYND domain-containing protein 4 | partial | Transcriptional repression through the recruitment of histone deacetylases. Immune response in drosophila. | no | no |
| 08F | 780576 | 780851 | SET and MYND domain-containing protein 4 | partial | Transcriptal repression through the recruitment of histone deacetylases. Immune response in drosophila. | no | yes |
| 08F | 789149 | 790123 | Uncharacterized protein | partial | Unknown | no | yes |
| 08F | 783612 | 787685 | Serine/threonine-protein phosphatase 2A 55 kDa regulatory subunit B | whole | Modulate substrate selectivity and catalytic activity. | yes | no |
| 08F | 786411 | 786753 | Major facilitator superfamily transporter | partial | Membrane transport proteins that facilitate movement of small solutes across cell membranes. | no | no |
| 08F | 874270 | 879847 | Beta-1,4-glucuronyltransferase 1 | whole | This enzyme is a type II transmembrane protein. It is essential for the synthesis of poly- N acetyl-lactosamine, a determinant for the blood group i antigen. | yes | yes |
| 08F | 867747 | 872574 | Uncharacterized protein | partial | Unknown | no | yes |
| 09F | 773493 | 776776 | Uncharacterized protein | partial | Unknown | no | yes |
| 09F | 776947 | 780230 | Non-genic | - | Unknown | no | no |
| 09F | 781015 | 789914 | A disintegrin and metalloproteinase with thrombospondin motifs 3 | whole | Zinc binding: down-regulation of cell proliferation | yes | yes |
| 11F | 1054965 | 1066483 | Mitogen-activated protein kinase organizer 1 | fraction | Molecular scaffold that directly binds prolyl hydroxylase domain 3 (PHD3), thereby increasing Hypoxia inducible factor-1 stability (HIF-1), which is involved in restraining inflammatory processes in *Drosophila* | yes | no |
| 14F | 2444324 | 2444682 | Uncharacterized protein | fraction | Unknown | no | no |
| 14F | 2446665 | 2449054 | Uncharacterized protein | whole | Unknown | no | yes |
| 14F | 2449538 | 2454549 | 4-hydroxyphenylacetate degradation bifunctional isomerase/decarboxylase | whole | This protein is involved in step 4 and 5 of the subpathway that synthesizes pyruvate and succinate semialdehyde from 4-hydroxyphenylacetate. | no | no |
| 14F | 185436 | 190137 | Epoxide hydrolase EPH1 protein | partial | Biotransformation enzyme that catalyzes the hydrolysis of arene and aliphatic epoxides to less reactive and more water soluble dihydrodiols by the trans addition of water. May play a role in the metabolism of endogenous lipids such as epoxide-containing fatty acids. | no | no |
| 23F | 467388 | 469341 | Non-genic | - | Unknown | no | no |
| 23F | 473200 | 475768 | F-box/LRR-repeat protein 14 | whole | Negative regulator of defense response among others. | no | yes |
| 26F | 1162330 | 1169622 | ABC protein, subfamily ABCH | whole | Transportation of a large diversity of substrates across lipid membranes. ABCH is found only in insects, which known lineage-specific expansions in *D. pulex*. | no | yes |
| 59F | 229898 | 232451 | Diacylglycerol kinase beta | whole | Induction of the T cell anergy. | no | no |
| 59F | 230339 | 232054 | Uncharacterized protein | partial | Unknown | no | yes |
| 59F | 233741 | 254611 | Eye-specific<br>diacylglycerol kinase | whole | Required for the maintenance<br>of phospholipid turnover within<br>the photoreceptor | no | no |

Four regions either have no genes, or they harbor immunity-unrelated genes or uncharacterized genes under purifying selection (Table 3 and 4). A possible explanation for this finding is that in these regions, it is not the genes themselves that are the target of selection, but rather non-genic sites such as transcription factors. Another possibility is that the gene annotations in these regions is either incomplete or incorrect.

In three candidate regions, we found uncharacterized genes with elevated ***π_N_/π_S_*** ratios. The high number of uncharacterized proteins is very common for species where functional gene annotation depends mainly on finding homologs to molecular model species such as *Drosophila melanogaster*. Further functional analysis will be necessary for understanding the role these regions play in the interaction of *D. magna* with the parasite *H. tvaerminnensis*, as well as other *Daphnia* parasites.

### Functional roles of candidate genes

As mentioned above there are five genes homologous to immunity related genes in other organisms. These genes might encode proteins involved in immunity even if the network to which they belong in *Daphnia* may not be the homologue of the one functioning in other organisms. The five genes are: a 1,4-beta Glucuronyl-transferase (GT), as well as four proteins known for their role in down regulating immune responses down regulation: Mitogen-activated protein kinase organizer 1 (MORG1), SET and MYND domain-containing protein 4 (Smyd), F-box protein and Diacylglycerol kinase beta (DGK). The role of GTs in immunity is not clear. In some beetle species, GTs are up-regulated during pathogen or immune-related oxidative stress (Lin et al., 2016). We find this protein encoded on contig 08F with elevated ***π_N_*** and no ***π_S_*** polymorphisms (Figure 3). If GTs are involved in the immune response of *D. magna* to *H. tvaerminnensis* as well, then they could plausibly be a likely target of diversifying selection. On contig 08F, we also find a Smyd protein (Figure 3). Despite other functions, Smyd proteins in *D. melanogaster* are known for repression of Toll-like-receptor 4 pathway through histone methylation (Calpena et al., 2015; Stender et al., 2012). Several proteins of the Smyd family are found in *D. magna* immune clusters. Signatures suggestive of positive selection, as well as high *F*_GT_ values, are found for this gene, indicating diversifying selection between the two population groups (Figure 3). The potential role in immune downregulation and the clear genetic summaries make this protein a likely target for diversifying selection in the metapopulation due to its role in host tolerance. On contig 11F, we find MORG-1, which is a well-known regulator of hypoxia-inducible factor (HIF) system, which, among others, is known to downregulate pro-inflammatory responses in *Drosophila* (Bandarra et al., 2015; Siddiq et al., 2007). We only find a fraction of the full protein, so it is possible that it is non-functional or has an alternate function at the given site, which could explain why it is under purifying selection (Figure 4). It could still have a central role in tolerance, especially because the introns seem to show very high *F*_GT_ values between population groups and it lies within the intervals of an earlier-described *H. tvaerminnensis* resistance QTL (Routtu & Ebert, 2015). The F-box protein is located on contig 23F (Figure 5) and inside the intervals of another QTL (Routtu & Ebert, 2015). This protein has many different functions but is found to play a role as a down regulator of plant resistance to fungi (Bonhomme et al., 2014), and it also is found to repress immune function through downregulation of a transcription factor in *D. melanogaster* (Khush et al., 2002; Muzzopappa & Wappner, 2005). The population genetic summaries around the gene are somewhat noisy, but we can still see a high *F*_GT_ within the gene, an elevated ***π_N_***, but no ***π_S_*** changes. Hence, if the F-box protein suppresses the immune response in *D. magna*, it is a likely target of increasing the level of tolerance and at the same time a likely target of diversifying selection (Figure 5).

**Figure 3.**
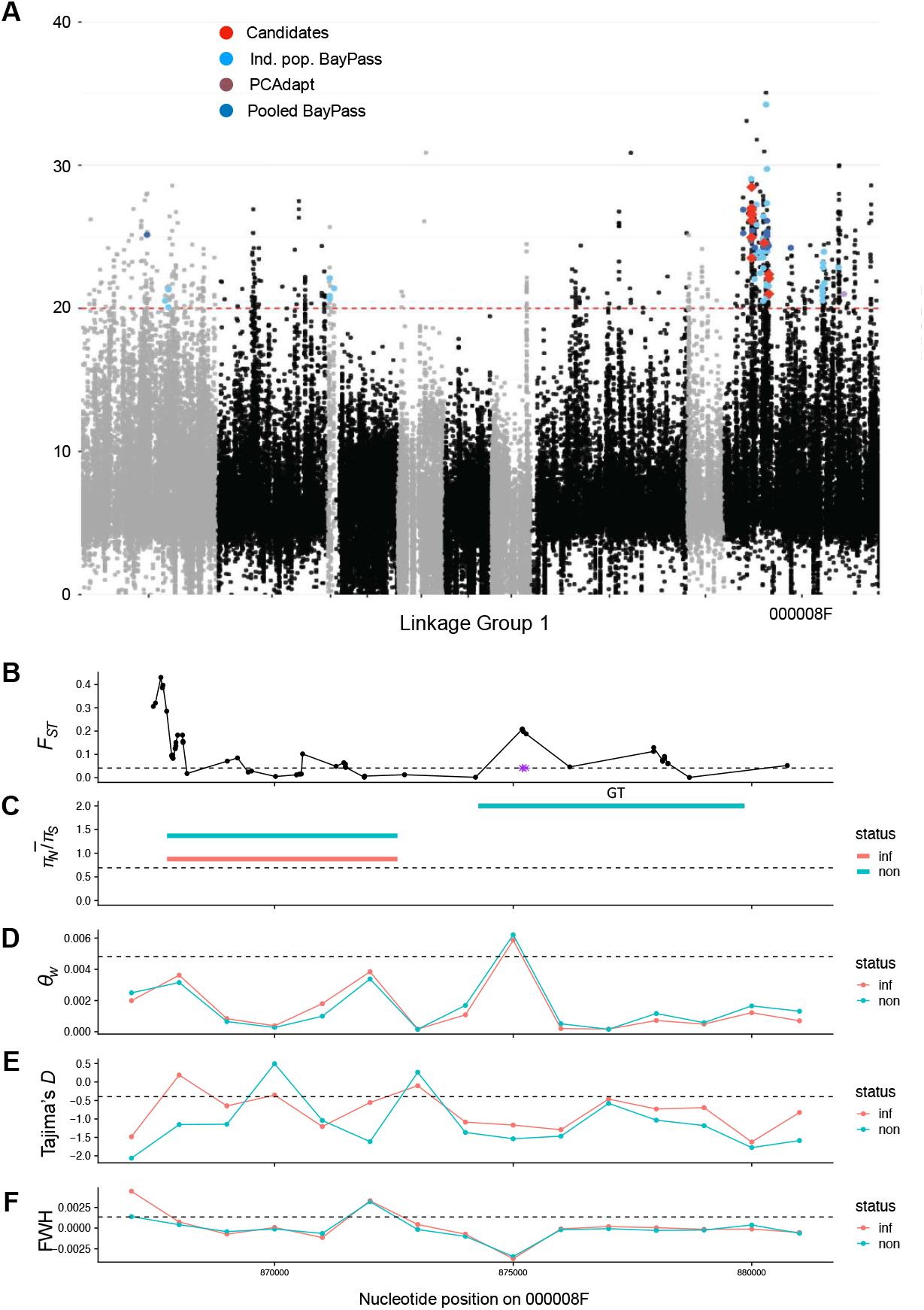
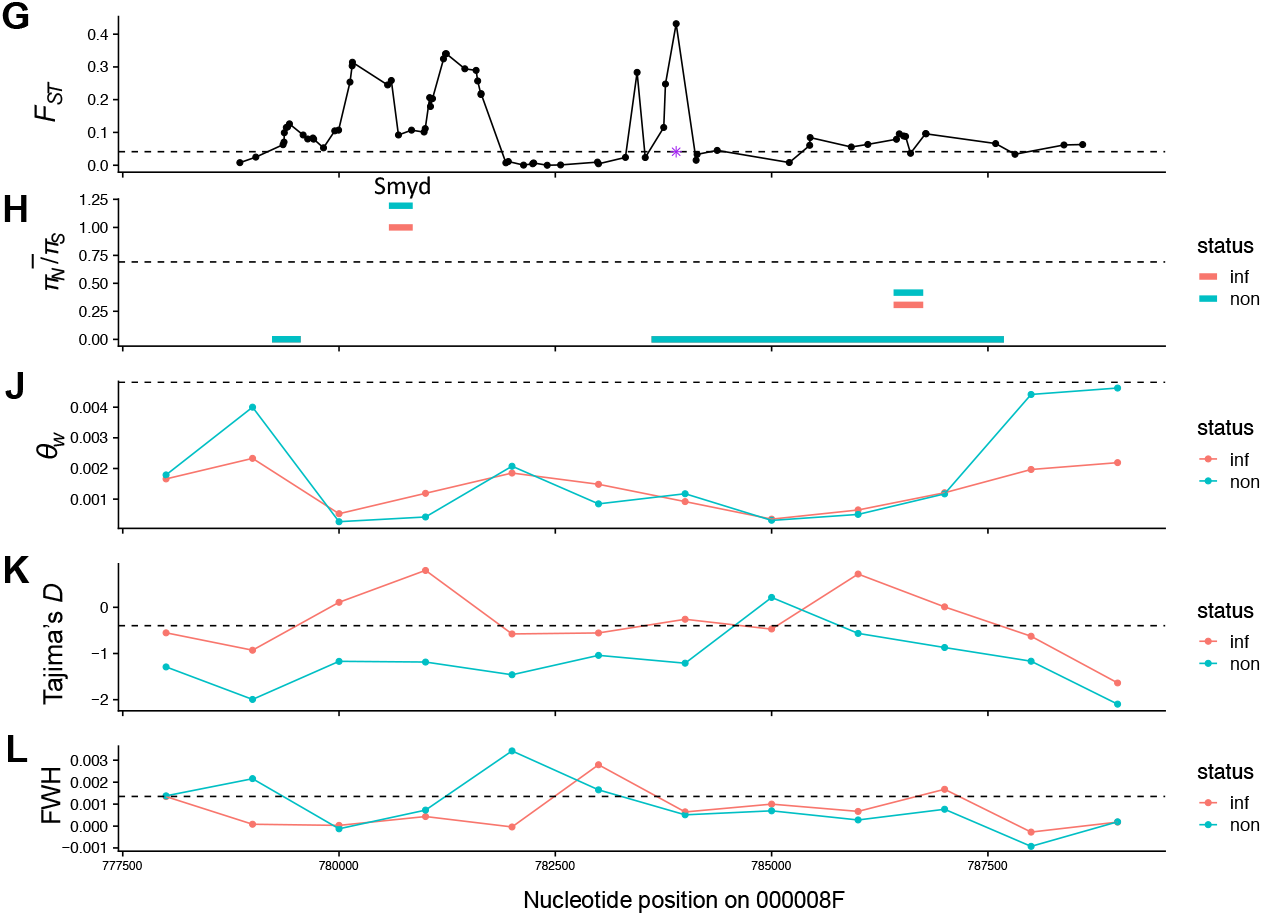
Evidence for selection on linkage group 1, contig 08F. A) The Bayes Factor (BF) for each SNP shows the degree of the association to the infection status on linkage group 1. Each point corresponds to a SNP. Gray and black are alternating for different contigs. The threshold line depicts 20 BF. Colored SNPs are outliers from different outlier tests and red SNPs (candidates) are outliers with signatures of positive selection. B-F) Population genetic summaries for candidate SNP and 12kb flanking region on different contigs. In purple is the candidate SNP highlighted. B and G) *F*_GT_ values of each SNP; C and H) mean ***π_N_/π_s_*** for each gene; D and J) genetic diversity; E and K) Tajima’s *D;* F and L) Fay & Wu’s H (FWH).

**Figure 4.**
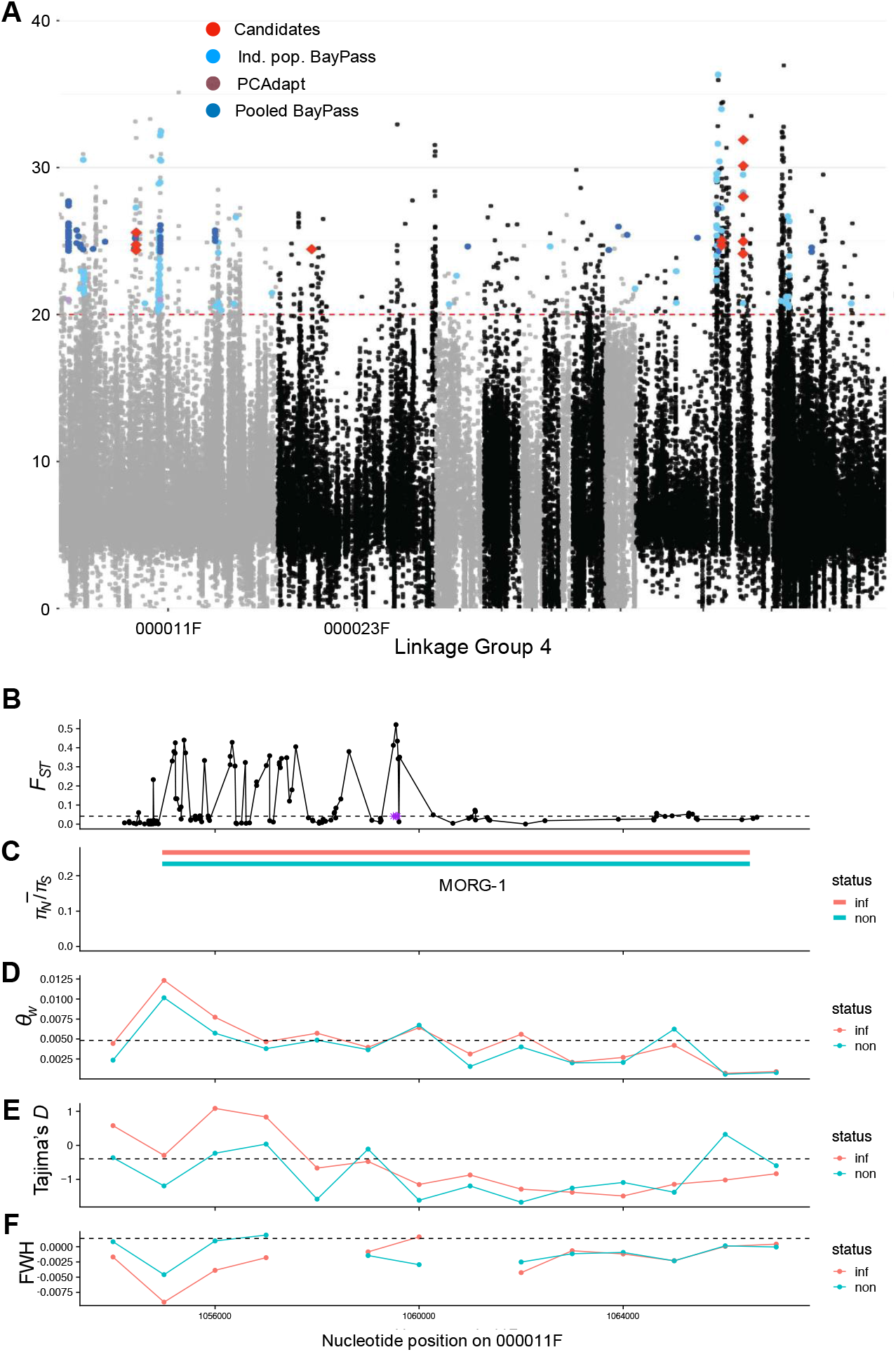
Evidence for selection on linkage group 4, contig 11F. A) The Bayes Factor (BF) for each SNP shows the degree of the association to the infection status on linkage group 4. Each point corresponds to a SNP. Gray and black are alternating for different contigs. The threshold line depicts 20 BF. Colored SNPs are outliers form different outlier-test and red SNPs (candidates) are outliers with signatures of positive selection. Population genetic summaries for candidate SNP and 12kb flanking region on different contigs. In purple is the candidate SNP highlighted. B) *F*_GT_ values of each SNP; C) mean ***π_N_/π_S_*** for each gene; D) genetic diversity; E) Tajima’s *D*; F) Fay & Wu’s H (FWH).

**Figure 5.**
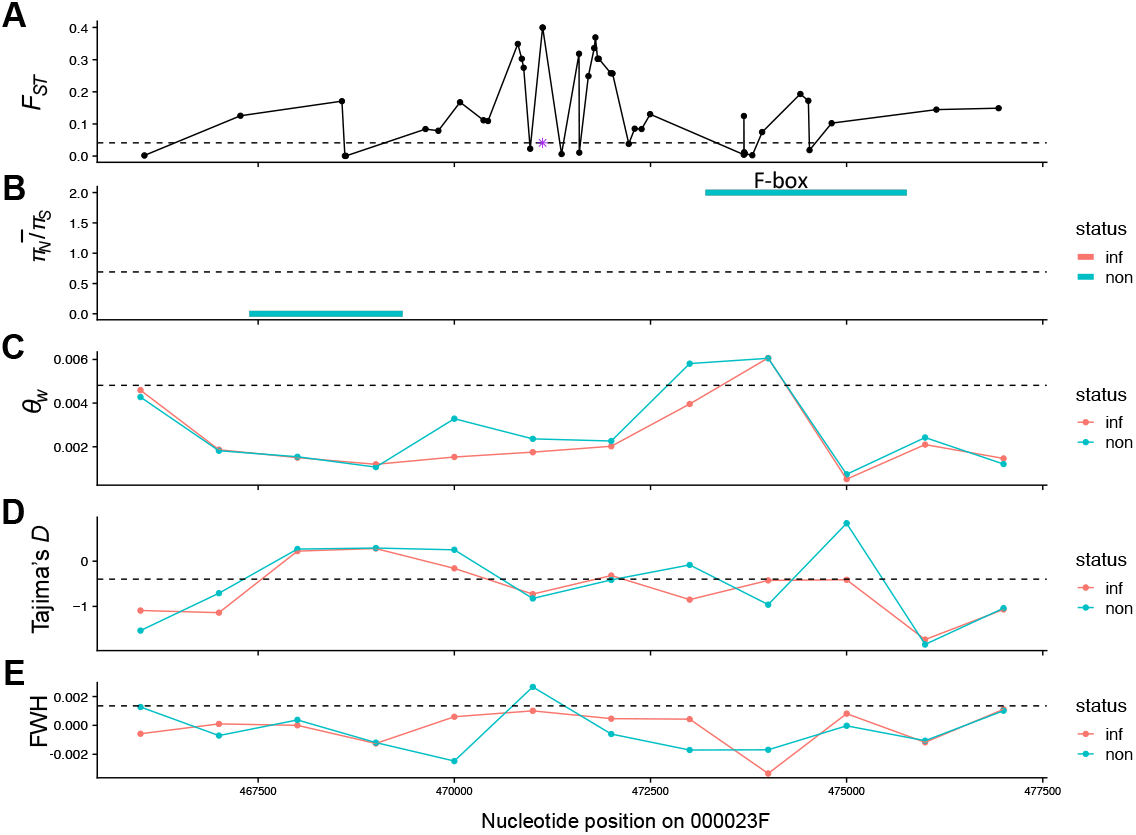
Evidence for selection on linkage group 4, contig 23F. In purple is the candidate SNP highlighted. A) *F*_GT_ values of each SNP; B) mean ***π_N_/π_s_*** for each gene; C) genetic diversity; D) Tajima’s *D*; E) Fay & Wu’s H (FWH).

Finally, we find a DGK on contig 59F (Figure 6). DGK belongs to a unique and conserved family of lipid kinases, providing a link between lipid metabolism and signaling in multicellular eukaryotes (Mérida et al., 2007). Especially in humans, these proteins are known for their prominent role as key negative regulators of immune function (Wattenberg & Raben, 2007). DGK seems to be under purifying selection on contig 59F in infected populations (Figure 7). If this gene is involved in *D. magna* downregulation of immune function, then the difference of the ***π_N_/π_S_*** rate we see in infected and non-infected populations could be because the gene is already in the state that increases tolerance.

**Figure 6.**
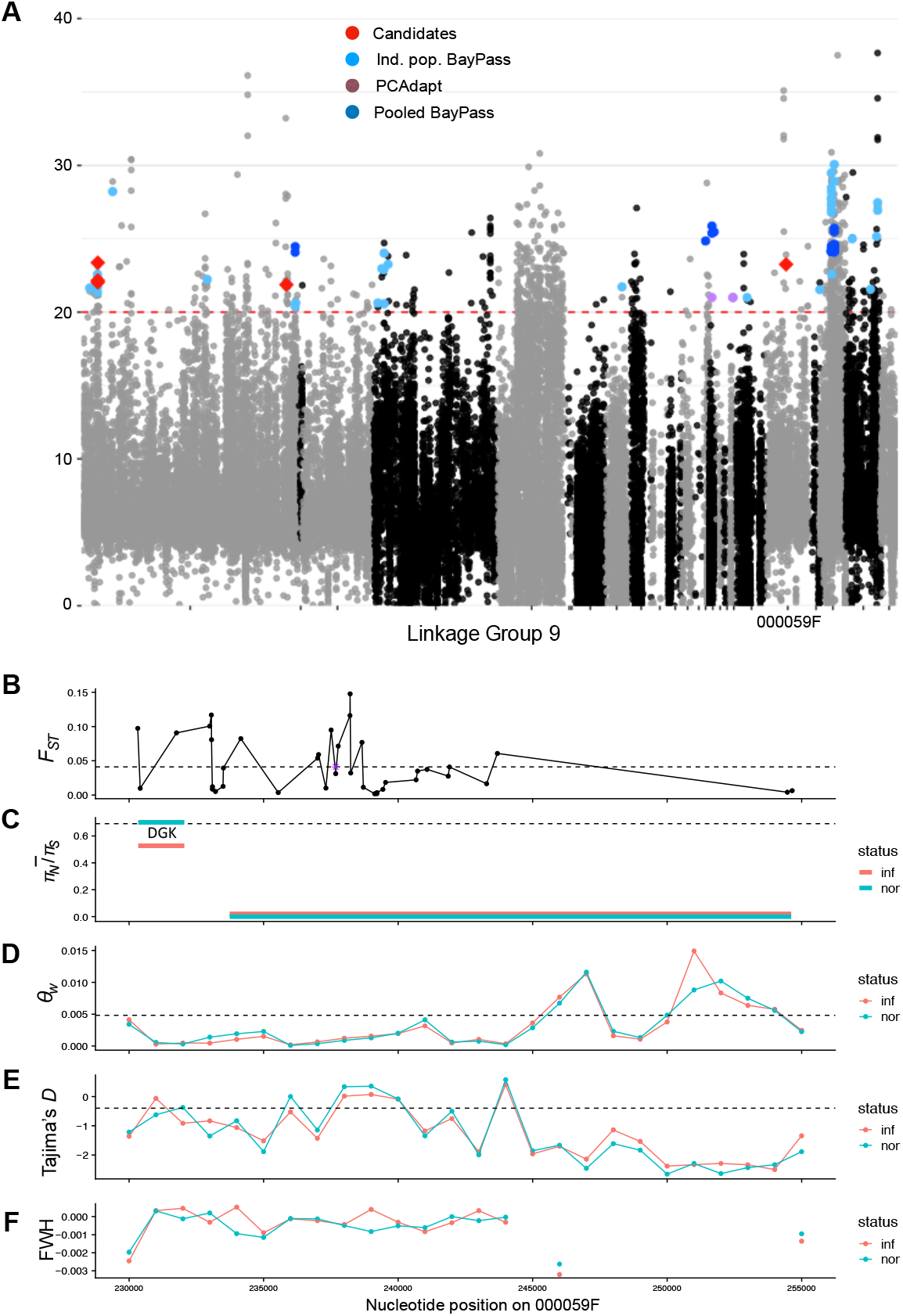
Evidence for selection on linkage group 9, contig 59F. A) The Bayes Factor (BF) for each SNP shows the degree of the association to the infection status on linkage group 9. Each point corresponds to a SNP. Gray and black are alternating for different contigs. The threshold line depicts 20 BF. Colored SNPs are outliers form different outlier-test and red SNPs (candidates) are outliers with signatures of positive selection. Population genetic summaries for candidate SNP and 12kb flanking region on different contigs. In purple is the candidate SNP highlighted. B) *F*_GT_ values of each SNP; C) mean ***π_N_/π_S_*** for each gene; D) genetic diversity; E) Tajima’s *D*; F) Fay & Wu’s H (FWH).

Taken together, we found five genes involved in immune function, with four of them being involved in immune downregulation in other species. As stated above, immune regulators are usually thought to show a trade-off between resistance and tolerance. In fact, Zbinden et al. (2008) found high mortality of individuals that evolved without the parasite once he exposed them to *H. tvaerminnensis*. This could be due to an up-regulated immune system, which “overshoots” and leads to damage in the host without being able to clear the infection (Schneider and Ayres 2008). This is detrimental for both the host and the parasite. The genes potentially involved in immune downregulation are therefore plausible candidate with trade-offs between absence and presence of *H. tvaerminnensis*, but empirical data of the protein and gene functions in *D. magna* are lacking.

### Tolerance alleles under diversifying selection

In only two of the 13 candidate regions, both infected and non-infected populations show significant signatures of positive selection (see Supplementary Table 3). But even if not significant in one of the population groups, the summaries statistics of the two groups are very similar. A possible explanation of this finding is that diversifying selection leads to shifts in allele frequencies in long-term infected versus non-infected populations. If the difference between the two subpopulations at these sites where entirely due to neutral processes, then we would expect to find signatures of positive selection only in subpopulations that had evolved with the parasite, while in subpopulations evolved without parasites, we would find more neutral summaries.

Maintenance of genetic variation in tolerance due to diversifying selection has been found in several plant and animal systems (Råberg et al, 2007; Read et al., 2008). Usually tolerance alleles are thought to go to fixation rather quickly, because tolerant hosts not only have the advantage of extended survival and/or fecundity when infected, but also have a competitive advantage over non-tolerant hosts by maintaining high parasite burden i.e., “the use of parasites as biological weapons” (Restif & Koella, 2004). But as soon as tolerance alleles are costly, they are expected to decrease in frequency once the infection is no longer present in the population (Horns & Hood, 2012).

Schneider and Ayres (2008) proposed two classes of tolerance mechanisms where resistance and tolerance have opposite linked effects and thus are predicted to show a trade-off. This first class of proteins are effectors that are important for immune response but at the same time can cause self-harm to the host and therefore reduce tolerance. An example is reactive oxygen species produced during immune response against a pathogen, but can at the same time cause severe pathology (Lambeth et al., 2007). A second class of proteins contains regulators, which control both resistance and tolerance. These regulators can selectively up-regulate various immune responses that fight infection, and therefore increase resistance, or alternatively induce damage to the host, and therefore decrease tolerance. Hence, such regulators are examples of factors that are predicted to show a trade-off between resistance and tolerance. A second class of proteins contains regulators which control both resistance and tolerance. These regulators can selectively up-regulate various immune responses that fight infection, and therefore increase resistance, and alternatively induce damage to the host, and therefore decrease tolerance. Hence, such regulators are example of factors that are predicted to show a tradeoff between resistance and tolerance. The involvement of such factors in the increased tolerance of adapted *D. magna* hosts could be a plausible explanation for the observed diversifying selection in the metapopulation. The tolerance alleles which down-regulate immune responses would be beneficial in the populations with *H. tvaerminnensis* infections, but when confronted with no or other pathogens, the same tolerance alleles could be problematic due to non-optimal immune regulation.

## Conclusions

By identifying genomic regions putatively involved in the adaptation of *D. magna* host tolerance and resistance, the current study contributes to the understanding of the underlying genetics of the interaction with the microsporidium *H. tvaerminnensis*. The overall genetic patterns we find in outlier SNPs match our expectation of several loci evolving under positive selection in the presence of the parasite. This is consistent with previous findings, where an ongoing arms race was suggested as the underlying genetic model of *D. magna* and *H. tvaerminnensis* interactions (Zbinden et al., 2008). With the current study, we further strengthened previous findings of a quantitative nature of the interaction in this system (Krebs et al., 2017; Routtu & Ebert, 2015; Routtu et al., 2014). While we do not know at this point how many loci are involved, we confirmed the roles of several previously described QTLs in modulating the coevolutionary process between *D. magna* and *H. tvaerminnensis* in the Finnish metapopulation. Taken together, this study contributes to the understanding of how diversifying selection can act in a metapopulation and expands the range of systems where knowledge of a quantitative trait of tolerance is found.

## Acknowledgements

We thank the following individuals for help with data collection in the field over the years of the survey: V.Ilmari Pajunen, Irmeli Pajunen, Jürgen Hottinger, Frida Ben-Ami, Andrea Cabalzar, Mikko Lehto, Jennifer Lohr, David Preiswerk, David Duneau, Katharina Ida Ebert, Gleb Georg Ebert, A. Marcelino, Y. Haag, E. Haag, C. Liautard-Haag, C. Haag, C. Reisser, C. Molinier, E. Hürlimann, and C. Mills. We would also like to thank members of the Ebert group very useful discussions. This work was supported by the Swiss National Science Foundation (SNSF; grant numbers 310030B, 166677, 310030, and 188887 to DE).

## Data accessibility

Analysis scripts are available at https://github.com/peterdfields/Halter_etal_2022. Raw data are available under NCBI SRA accessions X (Bioproject ID: X).

## Author contributions

All authors designed the study. MH and PDF analyzed the data and wrote the manuscript. All authors reviewed the manuscript.

