## Supplementary material for "A genomic analysis of parasite-mediated population differentiation in a metapopulation": Suppl Materials

Supplementary Materials

Supplementary Tables

Supplementary Table 1. Population genetic summaries with different window sizes and different flanking regions. Summaries of all 705 outlier SNP’s flanking regions and the flanking regions of 100,00 randomly drawn SNP’s. *p*-values of the Wilcoxon rank test between the two groups of each summary are reported for each summary. $\pi_N$ and $\pi_S$ as well as $d_N$ and $d_S$ are calculated only for polymorphisms inside genes intersecting the flanking regions. Non populations = non-infected populations; Inf populations = infected populations.

| | S | Watterson | Pi | Tajim's D | Fay & Wu's H | nonsyn. Pol. | syn. Pol. | nonsyn. Div. | syn. Div. | Fst |
|---|---|---|---|---|---|---|---|---|---|---|
| Outlier SNP's w1000 | 8.57921 | 0.00299 | 0.00274 | -0.53821 | 0.00099 | 0.11041 | 0.05012 | 1.03338 | 0.55948 | 0.0913 |
| Random SNP's w1000 | 13.18618 | 0.00481 | 0.00441 | -0.39647 | 0.00135 | 0.0925 | 0.04842 | 0.78524 | 0.45392 | 0.04117 |
| Pvalue w1000 | 0 | 0 | 0 | 0 | 0.00196 | 0 | 0 | 0 | 0 | 0 |
| Outlier SNP's w2000 | 17.07211 | 0.003 | 0.00274 | -0.65356 | 0.00101 | 0.21799 | 0.09862 | 2.02211 | 1.09554 | 0.0913 |
| Random SNP's w2000 | 26.20827 | 0.00482 | 0.00439 | -0.49963 | 0.00147 | 0.18342 | 0.09573 | 1.5525 | 0.89692 | 0.04117 |
| Pvalue w2000 | 0 | 0 | 0 | 0 | 0.02917 | 0 | 0 | 0 | 0 | 0 |
| Outlier SNP's w500 | 4.2792 | 0.003 | 0.00274 | -0.42581 | 0.00096 | 0.0561 | 0.02543 | 0.51482 | 0.28055 | 0.0913 |
| Random SNP's w500 | 6.59667 | 0.00479 | 0.00442 | -0.29391 | 0.00121 | 0.04612 | 0.02416 | 0.39173 | 0.22661 | 0.04117 |
| Pvalue w500 | 0 | 0 | 0 | 0 | 0.00801 | 0 | 0 | 0 | 0 | 0 |
| Outlier SNP's 20kb | 8.45804 | 0.00297 | 0.00269 | -0.56537 | 0.00089 | 0.08878 | 0.04559 | 0.89847 | 0.51826 | 0.08147 |
| Random SNP's 20kb | 12.5477 | 0.00464 | 0.00417 | -0.44137 | 0.00123 | 0.08436 | 0.04481 | 0.76643 | 0.4478 | 0.04163 |
| Pvalue 20kb | 0 | 0 | 0 | 0 | 0 | 0 | 0 | 0 | 0 | 0 |

Supplementary Table 2. List of all known QTLs for resistance of the *Daphnia magna* system.

| | | | Assembly 2.3 | | Assembly 2.4 | | PacBio Assembly | | | |
|---|---|---|---|---|---|---|---|---|---|---|
| Paper | Trait | Type of evidence | Marker position | lg (2.3) | Scaffold | position | PacBio contig | Position | lg (PB) | Side of lg |
| Bourgeois | Pasteuria | GWAS * | - | - | scaffold00568 | 38683-3 | 000016F | ~ 2019389 | 1 | Side A |
| Krebs | short & long persistance | QTL** | sc01302_1996 | 1 | | | 000016F | 2199288 | 1 | Side A |
| Luijckx | Ordospora | QTL | sc01302_1996 | 1 | | | 000016F | 2199288 | 1 | Side A |
| Krebs | long persistance | QTL | sc03191_677 | 1 | | | 000016F | 1589168 | 1 | Side A |
| Bourgeois | Pasteuria | GWAS | - | - | scaffold00512 | 259572- | 000008F | ~ 2758815 | 1 | Side B |
| Bourgeois | Pasteuria | GWAS | - | - | scaffold00868 | 655900- | 000000F | ~ 6096384 | 2 | Side B |
| Bourgeois | Pasteuria | GWAS | - | 3 | scaffold00626 | 408382- | 000011F | ~ 320931 | 4 | Side A |
| Routtu*** | horizontal infection (3B) | QTL | sc02114_353 | 3 | | | 000011F | 1161997 | 4 | Side A |
| Routtu | Pasteuria (ABC Locus) | QTL | sc00288_965 | 3 | | | 000011F | 2067352 | 4 | Side A |
| Bento | Pasteuria (ABC Locus) | Fine mapping | - | 3 | scaffold00944 | 1376677 | 000011F | ~ 2196556 | 4 | Side A |
| Routtu | horizontal infection (3A) | QTL | sc00773_975 | 3 | | | 000023F | 474442 | 4 | Side A |
| Krebs | short persistance | QTL | sc02593_1287 | 3 | | | 000026F | 962435 | 4 | Side B |
| Bourgeois | Pasteruia (C1 & C19) | GWAS | - | - | scaffold00687 | 271111- | 000018F | ~ 2292450 | 7 | Siede A |
| Luijckx | Ordospora | QTL | contig28778_1370 | 6 | | | 000005F | 1935968 | 8 | Side A |
| Routtu | vertical & horizontal infection | QTL | sc00606_2933 | 6 | | | 000005F | 1747463 | 8 | Side A |
| Krebs | long persistance | QTL | sc00815_3177 | 6 | | | 000031F | 1027112 | 8 | Side B |
| Luijckx | Ordospora | QTL | sc02738_155 | 7 | | | 000014F | 1463901 | 9 | Side A |
| Routtu | horizontal infection | QTL | contig14698_81 | 8 | | | 000006F | 2796755 | 10 | Side A |
| Bourgeois | Pasteuria | GWAS | - | - | scaffold01036 | 711675- | 000013F | ~ 1513535 | 10 | Side B |
| Luijckx | Ordospora | QTL | sc01110_4114 | 8 | | | 000013F | 2591884 | 10 | Side B |
| Routtu | horizontal infection | QTL | contig29113_349 | 8 | | | 000013F | 1442280 | 10 | Side B |

* Position information: Sequence of 50 bp in the middle of the associated area (reported is the first base)
** Position information: Marker SNP at highest peak on QTL map (2.3)
*** Paper states that marker SNPs of assembly 2.4 were used, but it was actually assembly 2.3

Supplementary Table 3. Outliers group into 13 regions. Except for two regions (09F and 11F) eighter summaries of infected populations or summaries of non-infected populations are significantly different from the flanking region of the random sample. *p*-values of the Wilcoxon rank test between the two groups of each summary are reported.

| Contig | Position (bp) | Window (bp) | Status | Fst | Pvalue Fst | S | Watterson | Pvalue Watterson | Pi | Pvalue Pi | Tajimas' D | Pvalue Tajima's D | Fay & Wu's_H | Pvalue Fay & Wu's H |
|---|---|---|---|---|---|---|---|---|---|---|---|---|---|---|
| 000001F | 1140938 | 1141500 | inf | 0.0662 | 0.0022 | 5.9444 | 0.0021 | 0.0005 | 0.0016 | 0 | -0.923 | 0.0004 | -0.0007 | 0 |
| 000001F | 1140938 | 1141500 | non | 0.0662 | 0.0022 | 6.7083 | 0.0023 | 0.0944 | 0.0016 | 0.0125 | -1.2154 | 0.0002 | -0.0002 | 0 |
| 000001F | 1140959 | 1141500 | inf | 0.0662 | 0.0022 | 5.9444 | 0.0021 | 0.0005 | 0.0016 | 0 | -0.923 | 0.0004 | -0.0007 | 0 |
| 000001F | 1140959 | 1141500 | non | 0.0662 | 0.0022 | 6.7083 | 0.0023 | 0.0944 | 0.0016 | 0.0125 | -1.2154 | 0.0002 | -0.0002 | 0 |
| 000005F | 2726439 | 2726500 | inf | 0.0769 | 0 | 2.875 | 0.001 | 0 | 0.0007 | 0 | -0.8854 | 0.0035 | -0.0001 | 0.0295 |
| 000005F | 2726439 | 2726500 | non | 0.0769 | 0 | 2.5417 | 0.0009 | 0 | 0.0006 | 0 | -1.1291 | 0.0065 | -0.0004 | 0.1893 |
| 000008F | 537496 | 537500 | inf | 0.24 | 0 | 6.1111 | 0.0022 | 0.0029 | 0.0018 | 0.0003 | -0.9358 | 0.0003 | -0.001 | 0 |
| 000008F | 537496 | 537500 | non | 0.24 | 0 | 6.4167 | 0.0022 | 0.0787 | 0.0016 | 0.0152 | -1.0329 | 0.0016 | -0.001 | 0.0009 |
| 000008F | 538810 | 539500 | inf | 0.232 | 0 | 7.1806 | 0.0026 | 0.0421 | 0.0021 | 0.0064 | -0.858 | 0.0019 | -0.0008 | 0.0005 |
| 000008F | 538810 | 539500 | non | 0.232 | 0 | 7.3333 | 0.0025 | 0.4258 | 0.0018 | 0.0875 | -1.1246 | 0.0002 | -0.0011 | 0.0005 |
| 000008F | 539073 | 539500 | inf | 0.232 | 0 | 7.1806 | 0.0026 | 0.0421 | 0.0021 | 0.0064 | -0.858 | 0.0019 | -0.0008 | 0.0005 |
| 000008F | 539073 | 539500 | non | 0.232 | 0 | 7.3333 | 0.0025 | 0.4258 | 0.0018 | 0.0875 | -1.1246 | 0.0002 | -0.0011 | 0.0005 |
| 000008F | 539076 | 539500 | inf | 0.232 | 0 | 7.1806 | 0.0026 | 0.0421 | 0.0021 | 0.0064 | -0.858 | 0.0019 | -0.0008 | 0.0005 |
| 000008F | 539076 | 539500 | non | 0.232 | 0 | 7.3333 | 0.0025 | 0.4258 | 0.0018 | 0.0875 | -1.1246 | 0.0002 | -0.0011 | 0.0005 |
| 000008F | 539105 | 539500 | inf | 0.232 | 0 | 7.1806 | 0.0026 | 0.0421 | 0.0021 | 0.0064 | -0.858 | 0.0019 | -0.0008 | 0.0005 |
| 000008F | 539105 | 539500 | non | 0.232 | 0 | 7.3333 | 0.0025 | 0.4258 | 0.0018 | 0.0875 | -1.1246 | 0.0002 | -0.0011 | 0.0005 |
| 000008F | 539141 | 539500 | inf | 0.232 | 0 | 7.1806 | 0.0026 | 0.0421 | 0.0021 | 0.0064 | -0.858 | 0.0019 | -0.0008 | 0.0005 |
| 000008F | 539141 | 539500 | non | 0.232 | 0 | 7.3333 | 0.0025 | 0.4258 | 0.0018 | 0.0875 | -1.1246 | 0.0002 | -0.0011 | 0.0005 |
| 000008F | 539175 | 539500 | inf | 0.232 | 0 | 7.1806 | 0.0026 | 0.0421 | 0.0021 | 0.0064 | -0.858 | 0.0019 | -0.0008 | 0.0005 |
| 000008F | 539175 | 539500 | non | 0.232 | 0 | 7.3333 | 0.0025 | 0.4258 | 0.0018 | 0.0875 | -1.1246 | 0.0002 | -0.0011 | 0.0005 |
| 000008F | 539568 | 540500 | inf | 0.2214 | 0 | 7.1389 | 0.0025 | 0.0285 | 0.0021 | 0.0051 | -0.8017 | 0.0047 | -0.0006 | 0.0011 |
| 000008F | 539568 | 540500 | non | 0.2214 | 0 | 7.2778 | 0.0024 | 0.2595 | 0.0017 | 0.0386 | -1.1422 | 0.0001 | -0.0009 | 0.002 |
| 000008F | 783897 | 784500 | inf | 0.1171 | 0 | 3.9583 | 0.0013 | 0 | 0.0012 | 0 | -0.4363 | 0.463 | 0.0008 | 0.0189 |
| 000008F | 783897 | 784500 | non | 0.1171 | 0 | 5.0833 | 0.0017 | 0.0003 | 0.0011 | 0 | -1.3257 | 0 | 0.0008 | 0.0161 |
| 000008F | 875187 | 875500 | inf | 0.0659 | 0.0289 | 4.1389 | 0.0014 | 0 | 0.001 | 0 | -0.8774 | 0.0038 | -0.0002 | 0.0903 |
| 000008F | 875187 | 875500 | non | 0.0659 | 0.0289 | 4.5972 | 0.0015 | 0 | 0.0009 | 0 | -1.2257 | 0.0002 | -0.0003 | 0.0387 |
| 000008F | 875195 | 875500 | inf | 0.0659 | 0.0289 | 4.1389 | 0.0014 | 0 | 0.001 | 0 | -0.8774 | 0.0038 | -0.0002 | 0.0903 |
| 000008F | 875195 | 875500 | non | 0.0659 | 0.0289 | 4.5972 | 0.0015 | 0 | 0.0009 | 0 | -1.2257 | 0.0002 | -0.0003 | 0.0387 |
| 000008F | 875273 | 875500 | inf | 0.0659 | 0.0289 | 4.1389 | 0.0014 | 0 | 0.001 | 0 | -0.8774 | 0.0038 | -0.0002 | 0.0903 |
| 000008F | 875273 | 875500 | non | 0.0659 | 0.0289 | 4.5972 | 0.0015 | 0 | 0.0009 | 0 | -1.2257 | 0.0002 | -0.0003 | 0.0387 |
| 000009F | 782166 | 782500 | inf | 0.0612 | 0.045 | 5.6667 | 0.0018 | 0.0001 | 0.0014 | 0 | -0.7683 | 0.0059 | 0.0014 | 0.0408 |
| 000009F | 782166 | 782500 | non | 0.0612 | 0.045 | 4.875 | 0.0016 | 0.0064 | 0.0011 | 0.0015 | -1.0753 | 0.0017 | 0.0009 | 0.0012 |
| 000011F | 1059495 | 1059500 | inf | 0.1014 | 0 | 13.8056 | 0.0052 | 0.0003 | 0.005 | 0.0023 | -0.5232 | 0.213 | -0.0024 | 0 |
| 000011F | 1059495 | 1059500 | non | 0.1014 | 0 | 12.4028 | 0.0044 | 0.0016 | 0.0036 | 0.0274 | -1.0241 | 0.0024 | -0.0012 | 0 |
| 000011F | 1059546 | 1060500 | inf | 0.1035 | 0.0002 | 13.2917 | 0.0049 | 0.0072 | 0.0046 | 0.0502 | -0.6462 | 0.0426 | -0.0022 | 0 |
| 000011F | 1059546 | 1060500 | non | 0.1035 | 0.0002 | 12.1389 | 0.0043 | 0.007 | 0.0034 | 0.0726 | -1.0131 | 0.0021 | -0.0013 | 0 |
| 000011F | 1059573 | 1060500 | inf | 0.1035 | 0.0002 | 13.2917 | 0.0049 | 0.0072 | 0.0046 | 0.0502 | -0.6462 | 0.0426 | -0.0022 | 0 |
| 000011F | 1059573 | 1060500 | non | 0.1035 | 0.0002 | 12.1389 | 0.0043 | 0.007 | 0.0034 | 0.0726 | -1.0131 | 0.0021 | -0.0013 | 0 |
| 000014F | 194952 | 195500 | inf | 0.0547 | 0.0115 | 7.0278 | 0.0026 | 0.0518 | 0.0022 | 0.0243 | -0.6535 | 0.0293 | 0.0002 | 0.0044 |
| 000014F | 194952 | 195500 | non | 0.0547 | 0.0115 | 5.9861 | 0.002 | 0.0423 | 0.0014 | 0.0026 | -1.3728 | 0 | -0.0005 | 0.0061 |
| 000014F | 194954 | 195500 | inf | 0.0547 | 0.0115 | 7.0278 | 0.0026 | 0.0518 | 0.0022 | 0.0243 | -0.6535 | 0.0293 | 0.0002 | 0.0044 |
| 000014F | 194954 | 195500 | non | 0.0547 | 0.0115 | 5.9861 | 0.002 | 0.0423 | 0.0014 | 0.0026 | -1.3728 | 0 | -0.0005 | 0.0061 |
| 000014F | 194955 | 195500 | inf | 0.0547 | 0.0115 | 7.0278 | 0.0026 | 0.0518 | 0.0022 | 0.0243 | -0.6535 | 0.0293 | 0.0002 | 0.0044 |
| 000014F | 194955 | 195500 | non | 0.0547 | 0.0115 | 5.9861 | 0.002 | 0.0423 | 0.0014 | 0.0026 | -1.3728 | 0 | -0.0005 | 0.0061 |
| 000023F | 471125 | 471500 | inf | 0.1332 | 0.0001 | 5.6944 | 0.0024 | 0.0133 | 0.002 | 0.0039 | -0.6576 | 0.041 | -0.0002 | 0.0229 |
| 000023F | 471125 | 471500 | non | 0.1332 | 0.0001 | 6.8472 | 0.0027 | 0.7177 | 0.0023 | 0.635 | -0.5534 | 0.8549 | -0.0003 | 0.0466 |
| 000026F | 1172840 | 1173500 | inf | 0.0895 | 0 | 10.5694 | 0.0038 | 0.2276 | 0.0033 | 0.4671 | -0.5858 | 0.1451 | -0.0005 | 0.0145 |
| 000026F | 1172840 | 1173500 | non | 0.0895 | 0 | 11.4583 | 0.0039 | 0.0019 | 0.003 | 0.0301 | -0.978 | 0.0025 | -0.0006 | 0 |
| 000026F | 1172850 | 1173500 | inf | 0.0895 | 0 | 10.5694 | 0.0038 | 0.2276 | 0.0033 | 0.4671 | -0.5858 | 0.1451 | -0.0005 | 0.0145 |
| 000026F | 1172850 | 1173500 | non | 0.0895 | 0 | 11.4583 | 0.0039 | 0.0019 | 0.003 | 0.0301 | -0.978 | 0.0025 | -0.0006 | 0 |
| 000026F | 1465000 | 1465500 | inf | 0.1082 | 0 | 7.7361 | 0.0036 | 0.8647 | 0.0032 | 0.9705 | -0.4778 | 0.3848 | -0.0012 | 0.0017 |
| 000026F | 1465000 | 1465500 | non | 0.1102 | 0 | 7.7917 | 0.0035 | 0.0147 | 0.0027 | 0.0935 | -1.0109 | 0.0045 | -0.0017 | 0 |
| 000026F | 1467342 | 1467500 | inf | 0.1102 | 0 | 8.3889 | 0.0037 | 0.6207 | 0.0034 | 0.504 | -0.2649 | 0.6947 | -0.0013 | 0.0009 |
| 000026F | 1467342 | 1467500 | non | 0.1102 | 0 | 8.0139 | 0.0034 | 0.0368 | 0.0028 | 0.1597 | -0.9496 | 0.015 | -0.0015 | 0 |
| 000026F | 1467468 | 1467500 | inf | 0.1064 | 0 | 8.3889 | 0.0037 | 0.6207 | 0.0034 | 0.504 | -0.2649 | 0.6947 | -0.0013 | 0.0009 |
| 000026F | 1467468 | 1467500 | non | 0.1082 | 0 | 8.0139 | 0.0034 | 0.0368 | 0.0028 | 0.1597 | -0.9496 | 0.015 | -0.0015 | 0 |
| 000026F | 1467497 | 1467500 | inf | 0.1102 | 0 | 8.3889 | 0.0037 | 0.6207 | 0.0034 | 0.504 | -0.2649 | 0.6947 | -0.0013 | 0.0009 |
| 000026F | 1467497 | 1467500 | non | 0.1102 | 0 | 8.0139 | 0.0034 | 0.0368 | 0.0028 | 0.1597 | -0.9496 | 0.015 | -0.0015 | 0 |
| 000026F | 1468100 | 1468500 | inf | 0.1102 | 0 | 8.3194 | 0.0037 | 0.6731 | 0.0034 | 0.6483 | -0.3452 | 0.8506 | -0.0013 | 0.0006 |
| 000026F | 1468100 | 1468500 | non | 0.1064 | 0 | 8.2222 | 0.0035 | 0.026 | 0.0028 | 0.1405 | -0.9678 | 0.0089 | -0.001 | 0.0001 |
| 000059F | 237681 | 238500 | inf | 0.0437 | 0.0424 | 3.2778 | 0.0011 | 0 | 0.0009 | 0 | -0.7337 | 0.0165 | -0.0002 | 0.0174 |
| 000059F | 237681 | 238500 | non | 0.0437 | 0.0424 | 3.9583 | 0.0013 | 0 | 0.001 | 0 | -0.7781 | 0.3252 | -0.0004 | 0.0104 |

Supplementary Figures

Supplementary Figure 1. Pooled sequencing pipeline. Summary of the pipeline used to detect variants from Illumina pooled-sequencing of *D. magna* rock pool populations.

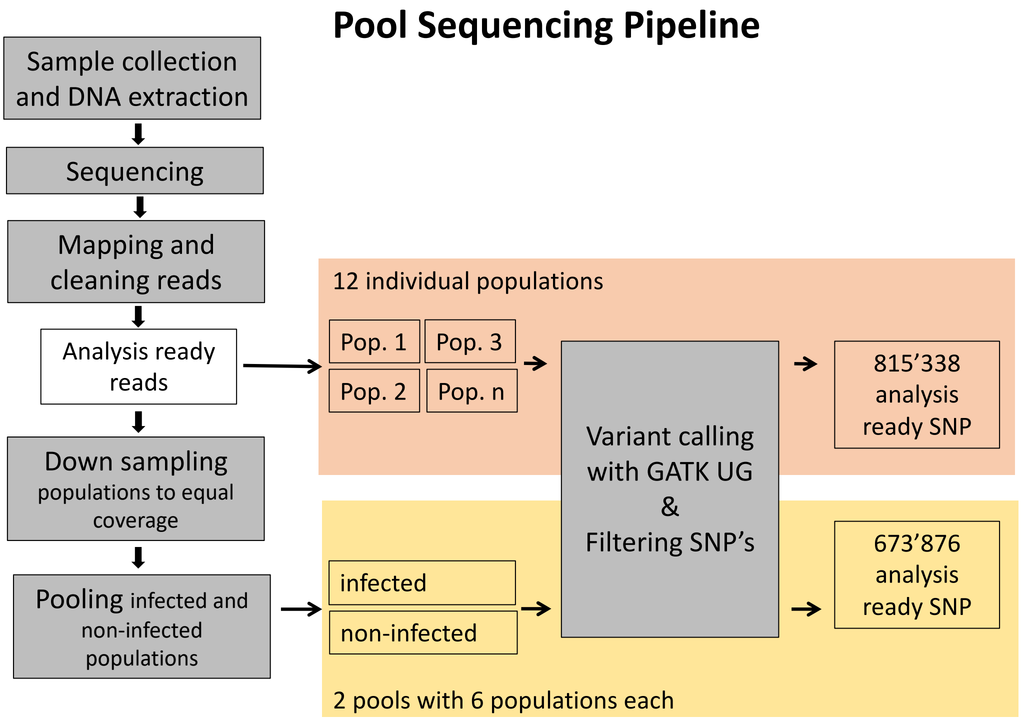

Supplementary Figure 2. Outlier analysis pipeline. Schematic of the pipeline used for outlier analysis and population genetic summaries used to find candidate genes.

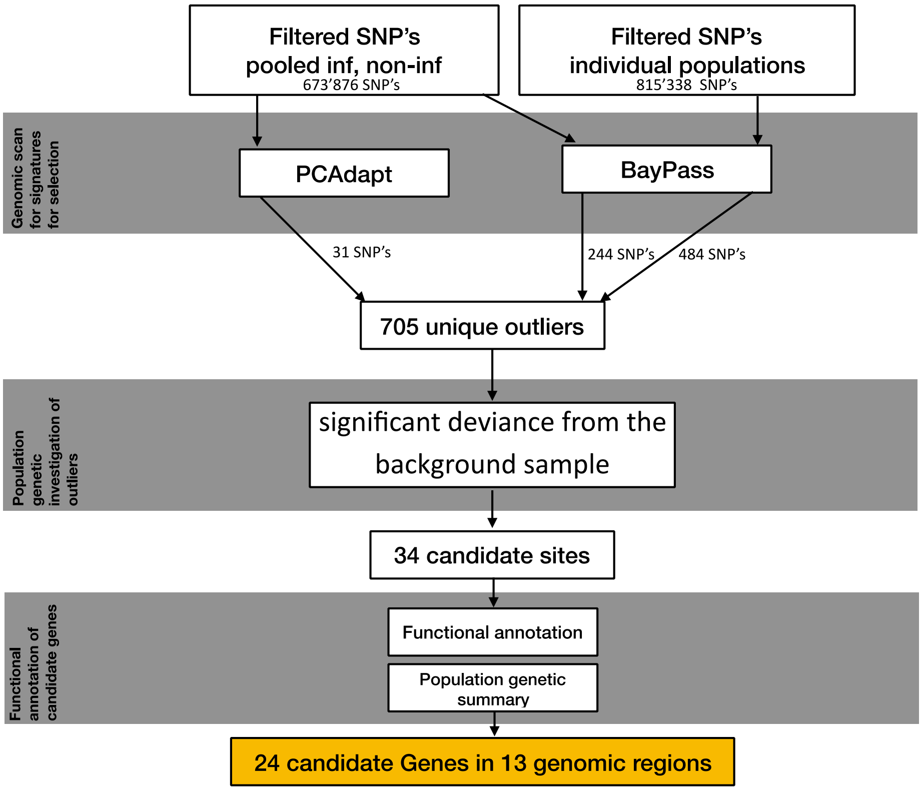

Supplementary Figure 3. Cumulative rank of all selection coefficients as estimated with SelEstim. Sites which were not identified as an outlier are labeled black while those which were identified as an outlier are labeled in red.

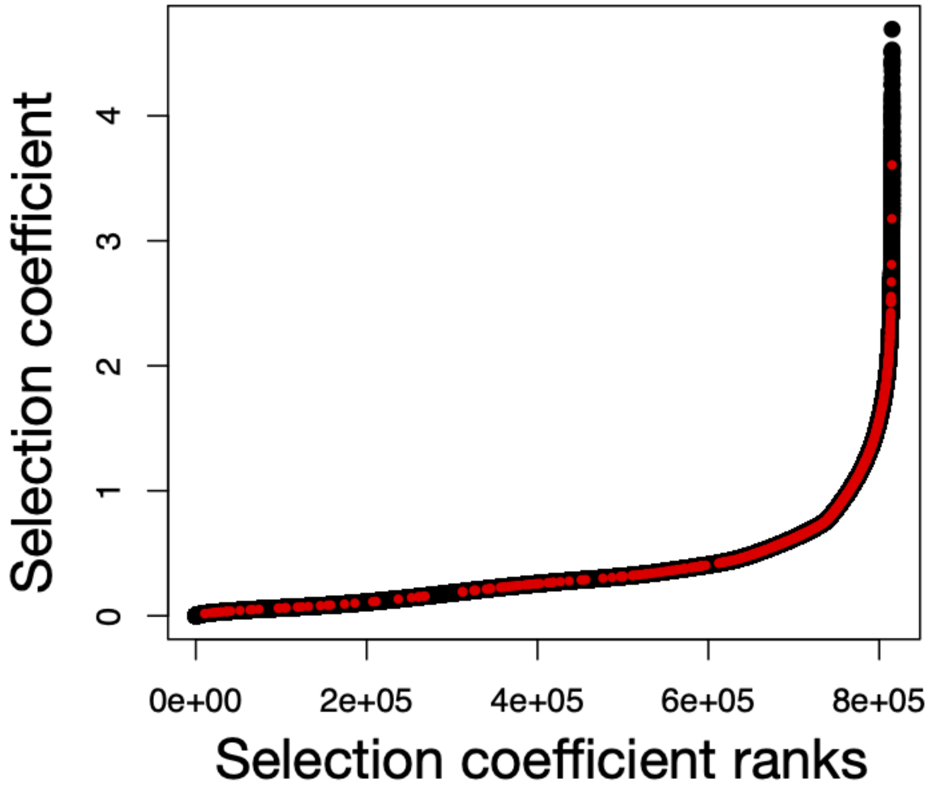

Supplementary Figure 4. The Bayes Factor (BF) for each SNP shows the degree of the association to the infection status on linkage group 1 – linkage group X. Each point corresponds to a SNP. Gray and black are alternating for different contigs. The threshold line depicts the 20 BF mark for decisive evidence of association. Colored SNPs are outliers form different outlier-test and red SNPs are outliers with signatures of positive selection. Linkage group X = contigs of unknown placement among the 10 linkage groups.

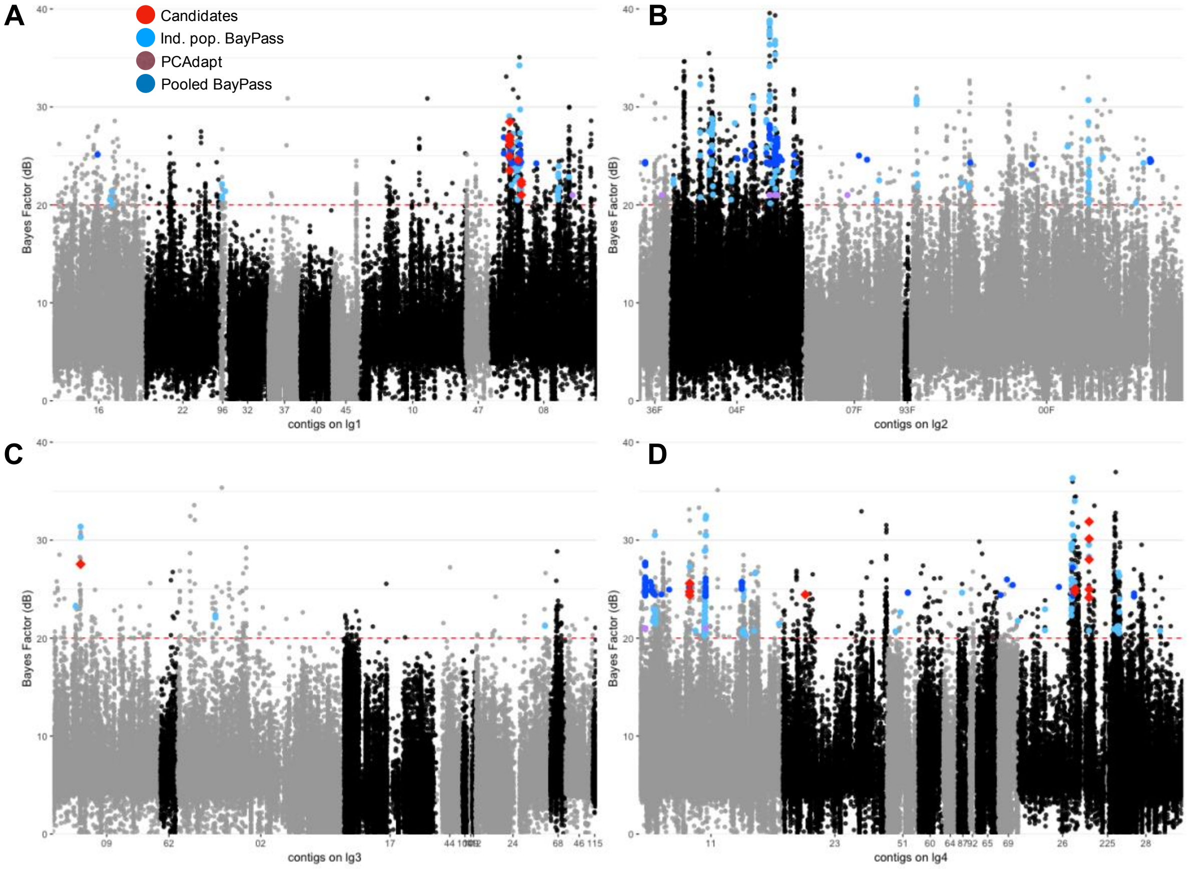

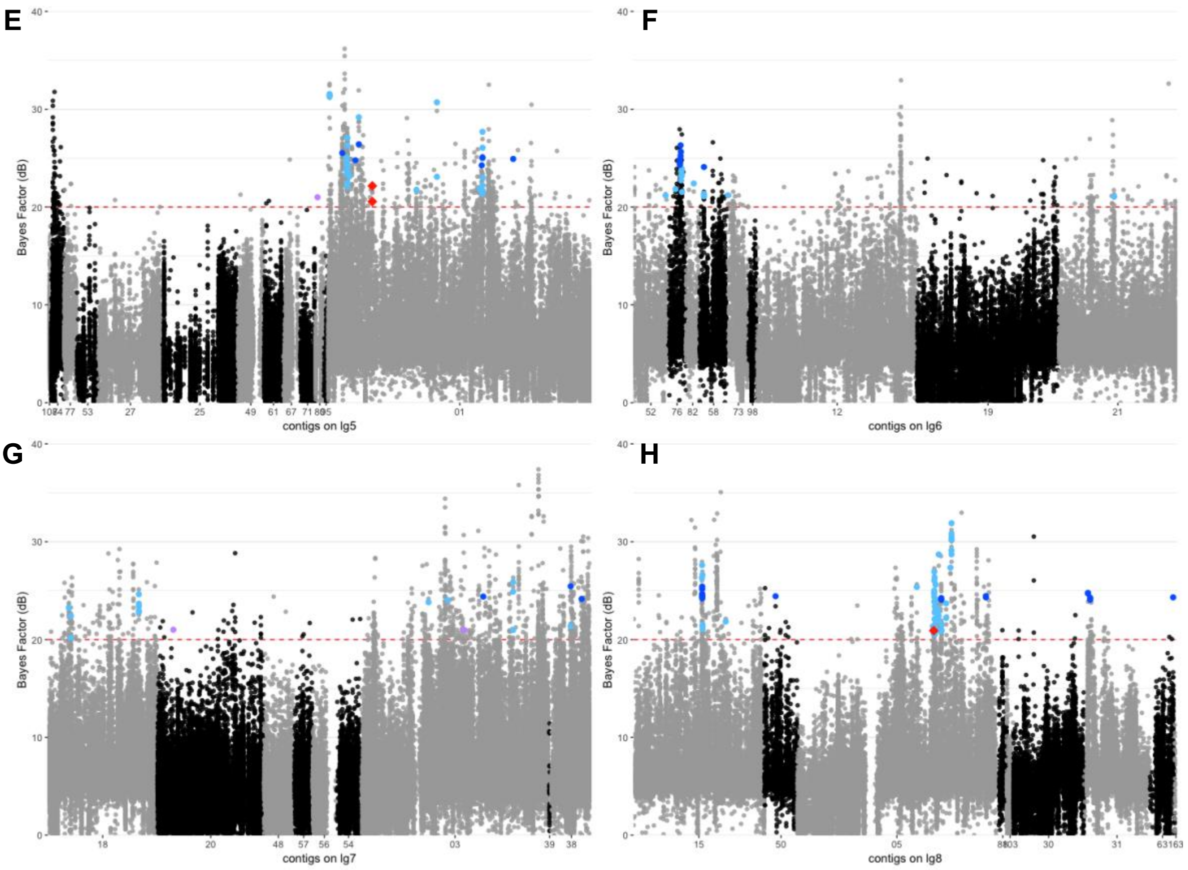
E
F
G
H
Bayes Factor (dB)
contigs on lg5
contigs on lg6
contigs on lg7
contigs on lg8

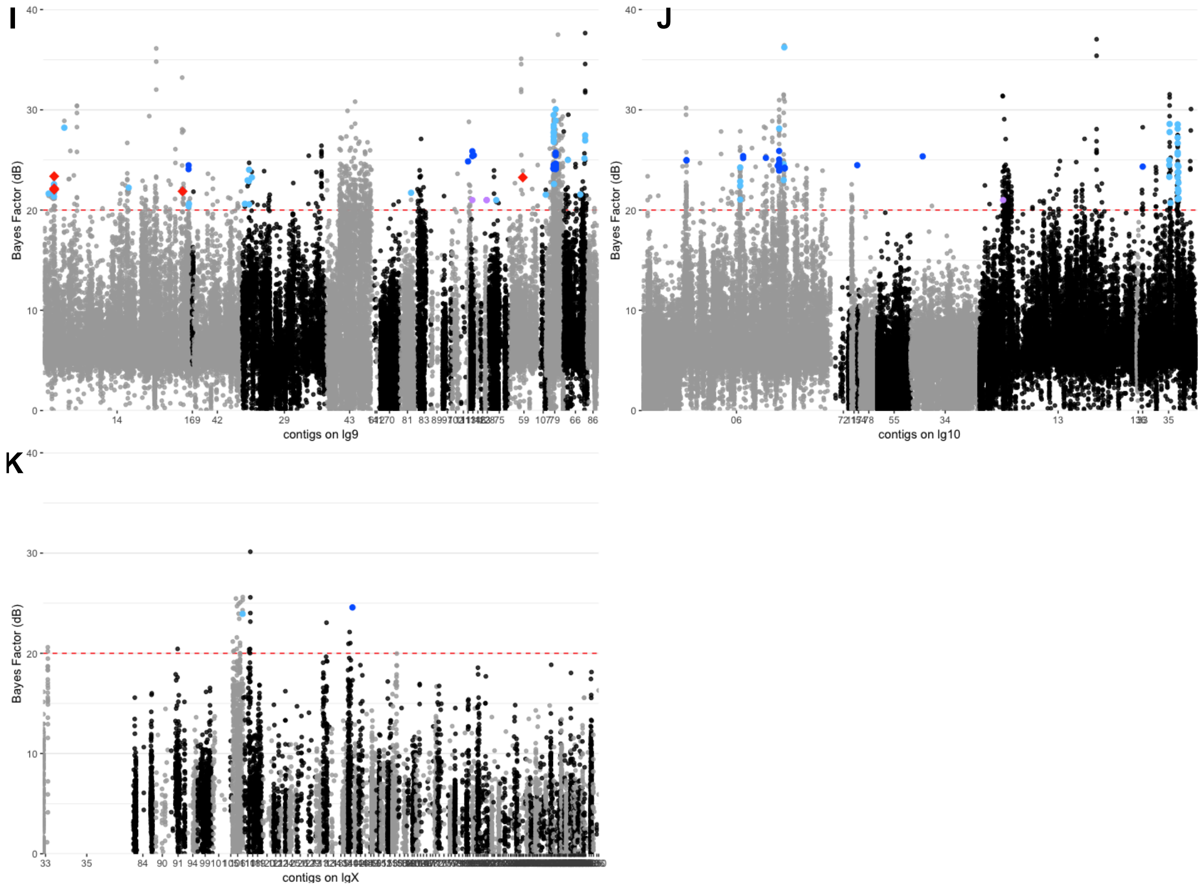
I
J
K
Bayes Factor (dB)
contigs on lg9
contigs on lg10
contigs on lgX
40
30
20
10
0
